# Ambisense transcription uncovers hidden coding capacity in compact circular RNA satellite viruses

**DOI:** 10.1101/2025.01.08.631871

**Authors:** Mai Kishimoto, Sakiho Imai, Akari Sono, Manabu Igarashi, Yukiko Sassa-O’Brien, Dennis Rubbenstroth, Masayuki Horie

## Abstract

Circular RNA replicons share compact, highly self-complementary genomes that impose severe constraints on genome organization and coding capacity. Recent studies have uncovered a broad diversity of such replicons, but their transcriptional and coding strategies remain inferred largely from sequence analyses. Here we show experimentally that some kolmioviruses (family *Kolmioviridae*), which have been described as negative-sense circular RNA satellite viruses, use ambisense transcription to expand coding capacity. We identify avian bornaviruses as helper viruses for Serinus canaria kolmiovirus (scKoV) and show that ambisense transcription drives expression of a previously unrecognized gene, open reading frame 2 (ORF2), that promotes helper-dependent transmission. Comparative analyses reveal ambisense gene expression in additional kolmiovirus lineages, suggesting independent acquisition during evolution. These findings provide experimental evidence that circular RNA replicons with highly constrained genomes can acquire additional protein-coding genes through ambisense transcription.

## 1. Introduction

Circular RNA replicons have long been exemplified by viroids and hepatitis delta virus (HDV; family *Kolmioviridae*), but recent studies have uncovered a much broader diversity of circular RNA elements, including ambiviruses, obelisks, and other proposed classes of viroid-like agents, such as epsilon and zeta viruses (*1–5*). Although these agents are highly diverse in sequence, many share a common genome architecture consisting of compact circular RNA genomes that form viroid-like, self-complementary, rod-like structures. Despite this broadly shared genome architecture, they are proposed to use diverse coding and gene expression strategies. However, for many of these recently identified circular RNA replicons, such strategies remain inferred primarily from sequence-based analyses and have yet to be tested experimentally.

Among such replicons, members of the family *Kolmioviridae* (KoVs) are small circular negative-sense RNA satellite viruses that depend on helper viruses for transmission. KoV genomes have been described as encoding only the delta antigen (DAg) and are highly self-complementary, viroid-like circular RNAs containing a promoter and catalytic ribozymes (*6*, *7*). Together, these features suggest that KoVs are under strong constraints on genome organization and coding capacity. In HDV, a representative virus in the family *Kolmioviridae*, however, coding capacity is expanded by host A-to-I RNA editing at the DAg stop codon, which generates two DAg isoforms with distinct roles in the viral life cycle: the small DAg isoform supports replication, whereas the large DAg isoform is required for virion assembly and transmission (*8–13*). By contrast, evidence that other KoVs, which were recently identified from vertebrates and invertebrates, use the same strategy remains limited (*14*), raising the possibility that they employ distinct gene expression strategies to expand coding capacity and regulate helper-dependent transmission.

Here we show that Serinus canaria kolmiovirus (scKoV; formerly referred to as Serinus canaria-associated deltavirus), which was identified in common canaries (*Serinus canaria*), encodes a novel protein-coding gene, designated open reading frame 2 (ORF2), on the genomic-sense strand and expresses it through ambisense gene expression. We further identify avian bornaviruses (family *Bornaviridae*) as helper viruses for scKoV and reveal that ORF2 contributes to helper virus-dependent transmission. Additionally, we show that ambisense gene expression also occurs in other KoV lineages, suggesting that this strategy may have been acquired independently during the evolution of KoVs. Together, these findings provide experimental evidence that circular RNA replicons can diversify their coding and gene expression strategies despite strong constraints on their genome architecture.

## 2. Results

### 2.1. Identification of the complete scKoV genome revealed ambisense gene expression and alternative splicing

scKoV is phylogenetically distinct from other mammalian and avian KoVs (Fig. 1A), and only its DAg-coding sequence had been identified (*15*). To gain further insights into the diversity of KoV genomes, we sought to determine the complete genome of scKoV by screening 15,452 publicly available bird RNA-seq datasets. Our mapping analysis identified 66 scKoV-positive datasets from four passerine and psittacine bird species, including one reported previously (*15*) (Table S1). From these datasets, we obtained 16 complete scKoV genome sequences across four BioProjects (Table S2 and Fig. S1). Phylogenetic analysis showed that these scKoVs are clustered into three distinct genotypes (Fig. S2A). Nucleotide sequence identities were >95% within each genotype and 83–89% between genotypes (Fig. S2B).

**Figure 1.**
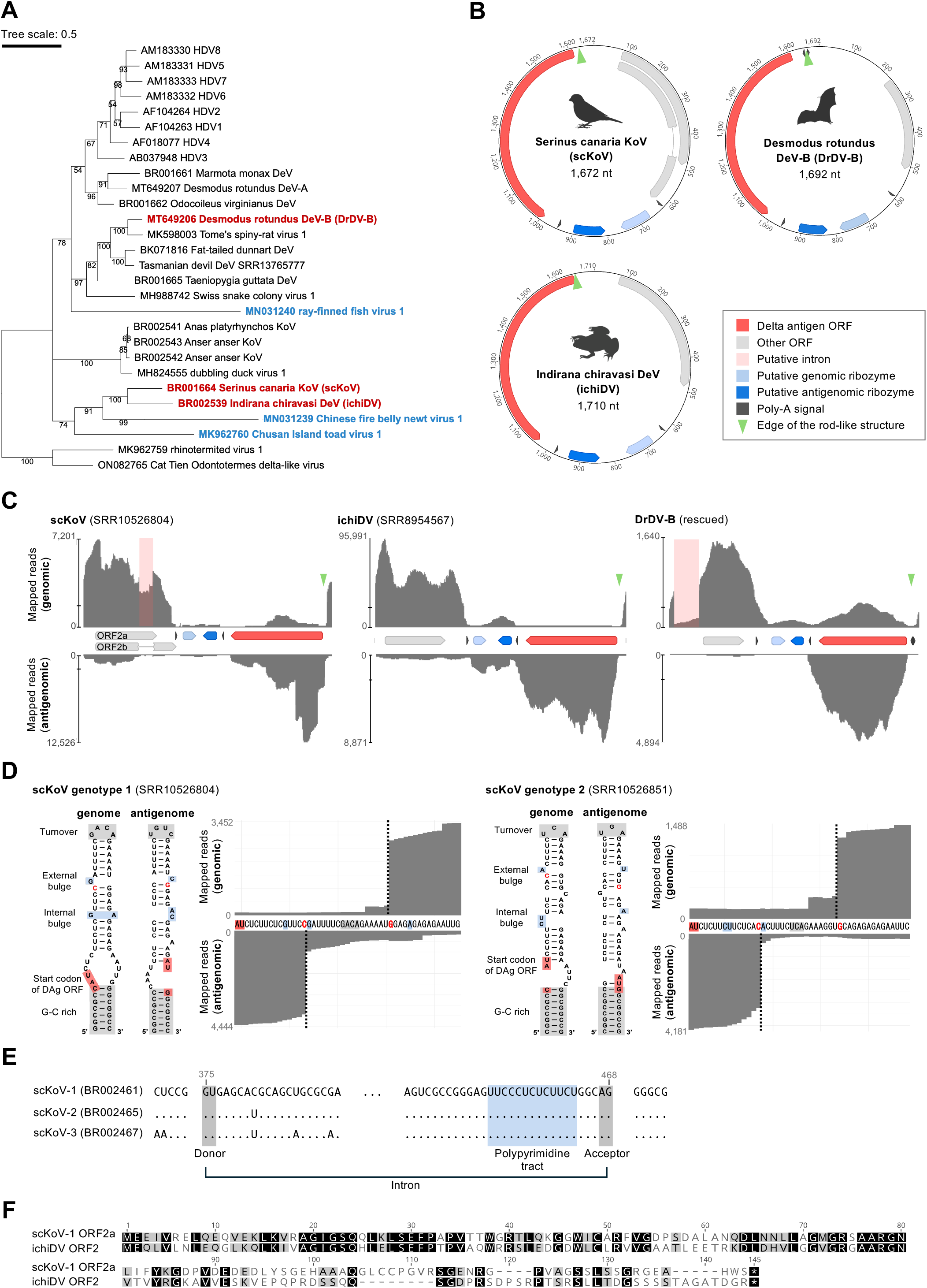
Identification of ambisense KoVs. (**A**) Phylogenetic tree of all known KoVs. The tree was inferred by the Bayesian Markov chain Monte Carlo method based on an amino acid sequence alignment of DAg. Posterior probabilities are shown for all nodes. KoVs with potential ambisense genome structures are shown in blue, and those with confirmed ambisense transcription patterns are shown in red. No GenBank accession number is available for Tasmanian devil deltavirus, as the sequence is not publicly available(*59*). The scale bar indicates the number of amino acid substitutions per site. (**B**) Genome organization of Serinus canaria KoV (scKoV), Indirana chiravasi deltavirus (ichiDV), and Desmodus rotundus deltavirus-B (DrDV-B). Colored arrows indicate annotations (ORFs, ribozymes, and poly-A signals). The numbers indicate nucleotide positions. The illustrations were created with BioRender.com. (**C**) Strand-specific mapping coverage of original short reads of scKoV, ichiDV, and DrDV-B. Colored arrows indicate annotations (ORFs, ribozymes, poly-A signals). Light pink box indicates putative intron. Green triangle indicates edge of the rod-like structure of circular genome. Information about other variants of scKoVs is provided in Fig. S5. (**D**) Secondary structures and expanded mapping patterns of the putative genomic and antigenomic promoter of scKoV. Conserved domains were annotated as previously reported(*22*). (**E**) Nucleotide alignment of predicted intron region using representative variants of scKoV in each genotype. Dots indicate common bases in the alignment. Gray and blue boxes indicate conserved GU-AG boundary and polypyrimidine tract, respectively. Gray numbers indicate nucleotide position in alignment of full-genome sequences. (**F**) Amino acid alignment of scKoV ORF2a and ichiDV ORF2. Identical amino acid residues are highlighted in black, similar substitutions in grey, and gaps are shown as dashes.

The scKoV genomes retain the characteristic features of KoV genomes, including lengths of 1,671–1,676 nucleotides (nt), GC contents of 54.0–55.4%, high self-complementarity, and a conserved genomic architecture containing the DAg ORF, poly-A signal, and ribozymes (Figs. 1B, S3, and S4) (*15–20*). The predicted ribozymes belong to the HDV class, with minor sequence variations that are unlikely to affect the secondary structures (Fig. S4) (*21*). Notably, an additional ORF, designated ORF2, together with a downstream poly-A signal, is present on the genomic-sense strand in the region complementary to the DAg ORF (Fig. 1B), and is strictly conserved in all sequences (Fig. S5), despite the genetic divergence of scKoV (Fig. S2). These findings suggest that scKoV encodes an additional protein-coding gene.

To assess this hypothesis, we mapped strand-specific mRNA-seq datasets (BioProject PRJNA591356 and PRJNA939570) to the corresponding viral genomes and examined the transcriptional pattern of scKoV. The mapping pattern suggested a genomic-sense transcript encoding ORF2 as well as the typical antigenomic-sense transcript encoding DAg (Figs. 1C and S5). Non-strand-specific datasets (PRJNA300534 and PRJNA679636) also showed a corresponding bimodal mapping pattern (Fig. S5C). The boundary between the genomic- and antigenomic-sense transcripts was predicted to be located at the rod edge of the scKoV genome, a region proposed to function as a transcriptional promoter (*22*). Furthermore, the putative transcription start sites of the genomic- and antigenomic-sense transcripts were predicted to be between the internal and external bulges at the rod edge (Fig. 1D), resembling HDV (*22*). The predicted secondary structure of the promoter was conserved across all sequences (Fig. S6). These results suggest that scKoV uses ambisense transcription.

This mapping analysis also detected splicing in the genomic-sense ORF2 transcript across all genotypes (Fig. 1C and S5). The putative intron retains the typical features of eukaryotic introns, including the GU–AG donor and acceptor motifs and a polypyrimidine tract upstream of the acceptor site (Fig. 1E) (*23*). Notably, the spliced ORF2 transcript could encode a protein (designated ORF2b) that differs from the protein encoded by the unspliced ORF2 transcript (designated ORF2a) in its C-terminal region (Fig. 1C and Table S3). These findings suggest that scKoV expresses genomic-sense transcripts that could encode two ORF2 isoforms.

HDV produces a larger DAg isoform through host A-to-I RNA editing at the stop codon of DAg ORF (*8–11*). To test whether scKoV uses the same mechanism, we examined nucleotide variations at the stop codon of DAg in the RNA-seq mapping data. Although putative A-to-I editing signatures (i.e., A-to-G substitutions) that would convert the DAg stop codon into a sense codon were detected in some datasets (Table S4), the predicted stop-codon readthrough would extend the DAg ORF beyond the downstream poly-A signal (Fig. S7), suggesting that scKoV is unlikely to express an alternative DAg isoform through host A-to-I RNA editing.

### 2.2. ORF2 was expressed during scKoV replication

To examine whether scKoV expresses ORF2 protein during replication, we attempted to rescue scKoV by transfecting a plasmid encoding a 1.2×-length scKoV genome into three avian cell lines (QT6, LMH, and DF-1), which can artificially initiate KoV replication cycle (*24*). By immunofluorescence assay (IFA), we detected DAg in all cell lines at 3 and 7 days post-transfection (dpt) (Figs. 2A and S8). Because the plasmid expresses genomic-sense scKoV RNA, from which DAg is not directly translated, detection of DAg, particularly at 7 dpt, indicates that scKoV was successfully rescued in cultured cells (*14*). Under the same conditions, we also detected ORF2-specific signals in all cell lines at 3 and 7 dpt (Figs. 2A and S8). Western blotting further detected a clear band for DAg and a single ORF2 band consistent with the predicted size of ORF2a (14.4 kDa), whereas no band corresponding to ORF2b (16.4 kDa) was observed (Fig. 2B). These results indicate that scKoV expresses ORF2 protein during replication.

**Figure 2.**
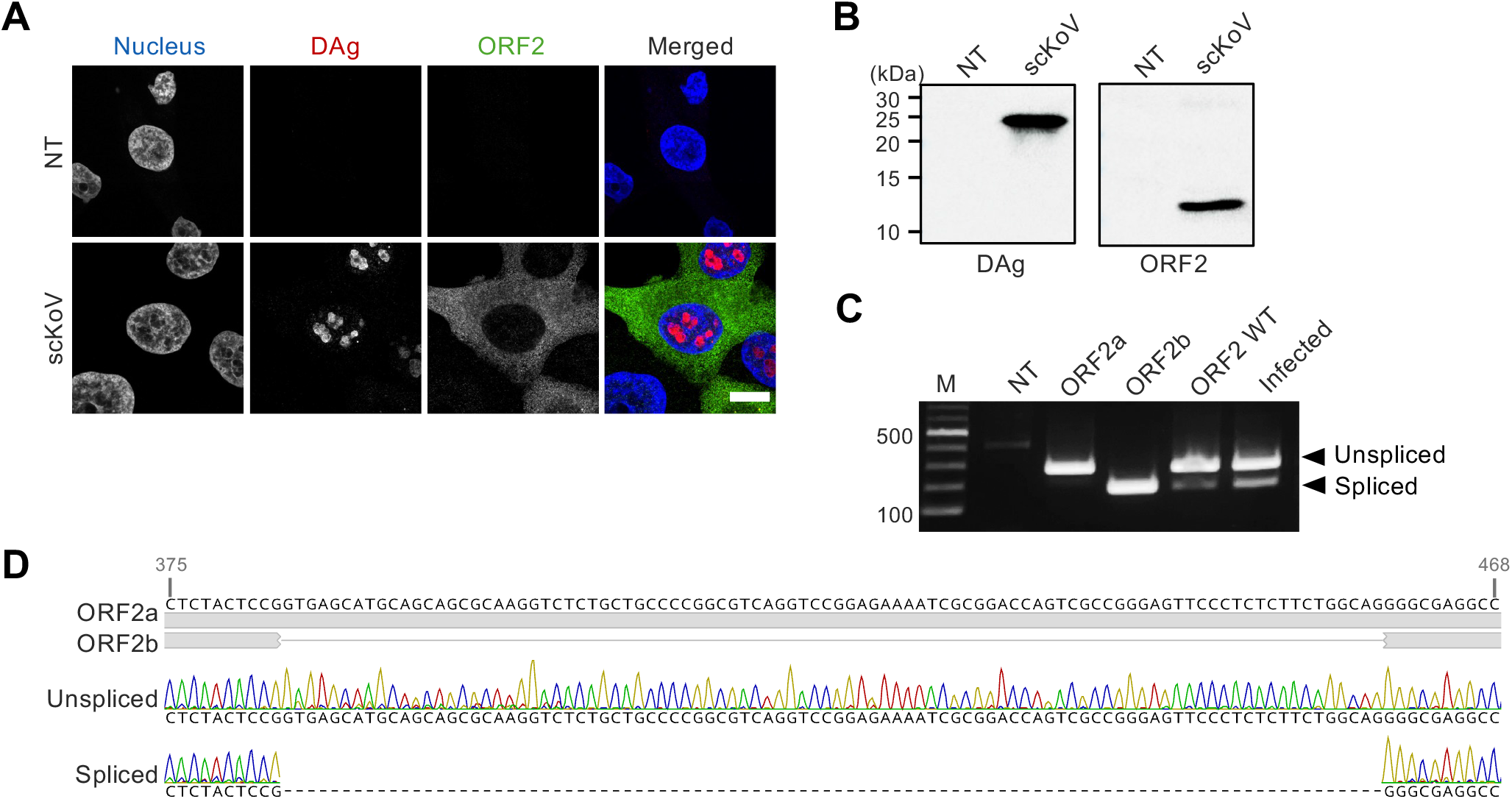
ORF2 expression during scKoV replication in cultured cells. (**A**) Subcellular localization of scKoV proteins in infected QT6 cells was analyzed by indirect immunofluorescence assay (IFA) using antibodies against scKoV DAg and ORF2. IFA images were acquired using confocal microscopy. Blue; DAPI, Red; DAg, Green; ORF2. Scale bars; 10 μm. (**B**) Western blotting analysis of scKoV-infected QT6 cells. The numbers indicate protein marker size (kDa). (**C**) RT-PCR to detect spliced and unspliced forms of scKoV ORF2 mRNA. M; 100 bp ladder. RNA from QT6 cells transfected with ORF2a, ORF2b, and ORF2 WT constructs (used in Figs. 3B and 3C) was used as size controls. (**D**) RT-PCR products of spliced and unspliced forms of scKoV ORF2 mRNA were validated using Sanger sequencing.

Because only a single ORF2 band was detected by western blotting, we next examined whether the scKoV ORF2 transcript undergoes alternative splicing in infected cells. RT-PCR across the predicted splice junction yielded amplicons corresponding to both spliced and unspliced transcripts, with the spliced transcript detected at a much lower level than the unspliced transcript (Figs. 2C and 2D). These results suggest that splicing of the ORF2 transcript occurs at low efficiency, which may explain the lack of detectable ORF2b protein expression.

### 2.3. ORF2 is dispensable for scKoV replication

To investigate the potential function of ORF2 proteins, we performed *in silico* analyses including sequence similarity searches, structural predictions, and functional domain searches (see Supporting text 1). However, these analyses did not identify any recognizable functional features or conserved domains (Fig. S9).

To experimentally investigate the function of ORF2, we assessed whether it is required for viral replication. To do so, we constructed a plasmid encoding a 1.2×-length scKoV genome with a mutation at the start codon of ORF2 (pscKoV-ΔORF2), together with a corresponding plasmid carrying a mutation at the DAg start codon (pscKoV-ΔDAg) as a control, and transfected them into cells (Fig. S10). In cells transfected with pscKoV-ΔDAg, as expected, neither DAg nor ORF2 protein expression was detected. In contrast, in cells transfected with pscKoV-ΔORF2, DAg was detected at 3 and 7 dpt (Fig. 3A), indicating that scKoV replicated in the absence of ORF2. These results demonstrate that ORF2 is dispensable for scKoV replication.

**Figure 3.**
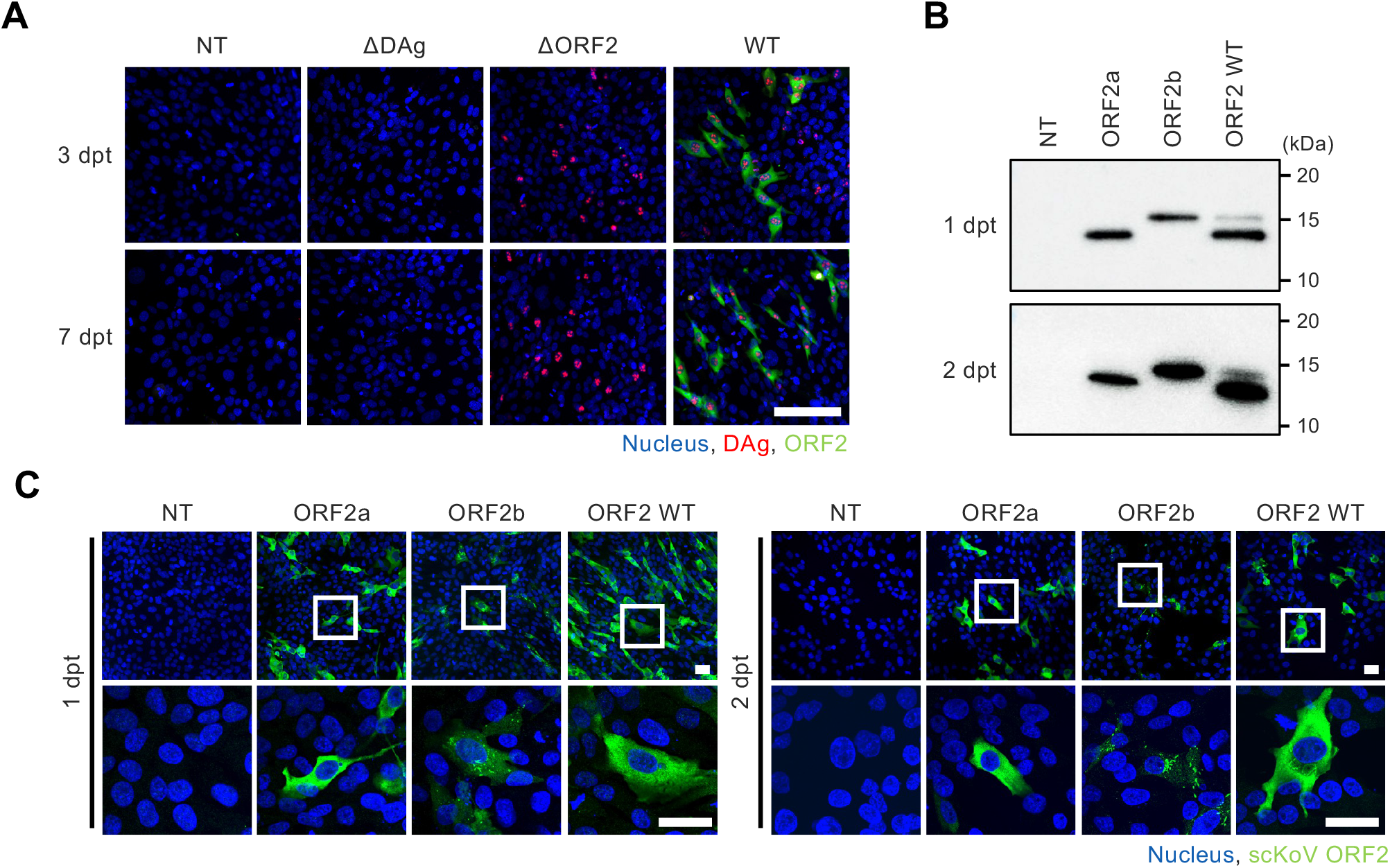
ORF2 is dispensable for scKoV replication and shows isoform-specific localization. (**A**) Indirect immunofluorescence assay (IFA) of QT6 cells transfected with plasmids encoding translation-deficient scKoV mutants. (**B and C**) QT6 cells transfected with plasmids expressing ORF2a, ORF2b, or ORF2 WT (see Fig. S11) were analyzed at 1 and 2 days post-transfection using western blotting (B) and IFA (C). Non-transfected cells (NT) were used as negative controls. IFA images were acquired using confocal microscopy. Blue; DAPI, Red; DAg, Green; ORF2. Scale bars; 100 μm (A), and 30 μm (C).

### 2.4. ORF2a and ORF2b show distinct subcellular localization

To gain insights into the function of ORF2, we examined the subcellular localization of scKoV proteins. In scKoV-infected cells, ORF2 proteins were diffusely distributed in the cytoplasm. By contrast, DAg formed irregularly shaped inclusion bodies in the nucleus that are characteristic of KoVs (Fig. 2A).

We next examined whether this cytoplasmic localization differed between the two ORF2 isoforms. We constructed plasmids expressing either ORF2a or ORF2b (Fig. S11). Western blotting confirmed expression of both proteins at predicted molecular weights, with cells expressing ORF2 WT showing two corresponding bands (Fig. 3B). Notably, in cells expressing ORF2 WT, ORF2b expression was markedly lower than that of ORF2a, which is consistent with infected cells. IFA revealed that both ORF2a and ORF2b proteins localized to the cytoplasm, but ORF2a was diffusely distributed, whereas ORF2b formed puncta (Fig. 3C). These ORF2b structures were more prominent at 2 dpt than at 1 dpt. The localization pattern of ORF2 WT resembled that of ORF2a, likely reflecting its higher expression level.

### 2.5. Avian bornaviruses are potential helper viruses for scKoV

To investigate whether ORF2 contributes to the transmission of scKoV, we first searched for potential helper viruses in scKoV-positive RNA-seq datasets. BLAST searches using the assembled contigs obtained from scKoV-positive RNA-seq datasets identified avian bornaviruses (genus *Orthobornavirus* of the family *Bornaviridae*) in 64 of 66 datasets (Figs. 4A and S12): canary bornavirus 1 and 3 (CnBV-1 and 3; *Orthobornavirus serini*) were found in datasets from *S. canaria*, *S. serinus*, and *H. mexicanus* (order Passeriformes), whereas parrot bornavirus 2 (PaBV-2; *Orthobornavirus alphapsittaciforme*) was present in datasets from *M. monachus* (order Psittaciformes). No other enveloped viruses were detected. These results suggest that orthobornaviruses are candidates for scKoV helper viruses.

**Figure 4.**
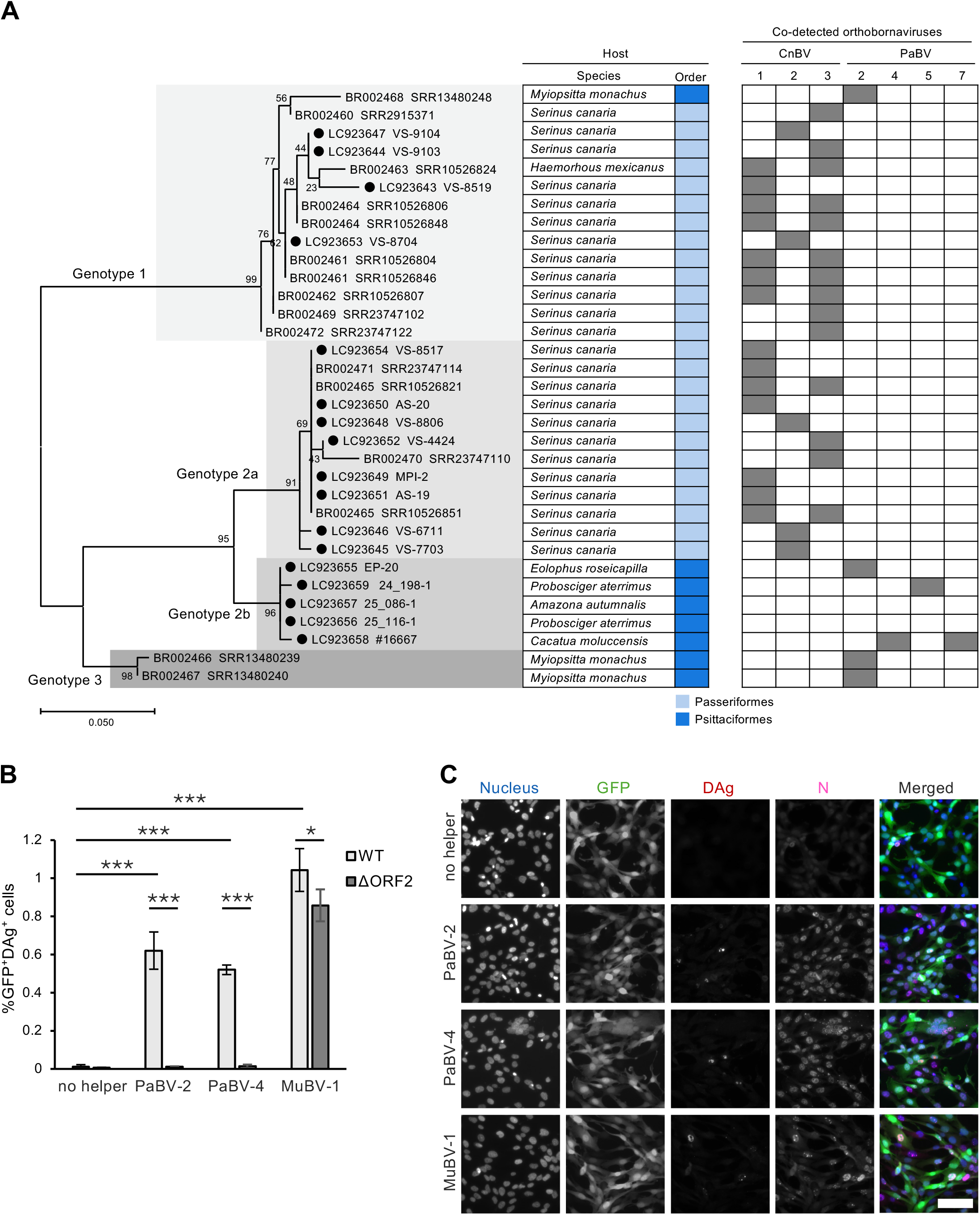
Avian bornaviruses support scKoV transmission. (**A**) Phylogenetic tree of partial scKoV DAg nucleotide sequences (197 bp; genome positions 1,116 to 1,312) detected in public RNA-seq datasets and diagnostic bird samples (indicated by black dots). Host species and co-detected canary bornaviruses (CnBVs) and parrot bornaviruses (PaBVs) are indicated for each dataset or sample (indicated by grey boxes). Only RNA-seq datasets yielding complete scKoV genomes are shown here; full information of public RNA-seq data is provided in Fig. S12. (**B**) Transmission efficiency of scKoV wild type (WT) or ORF2-deficient mutant (ΔORF2) in the presence or absence of PaBV-2, PaBV-4, or munia bornavirus 1 (MuBV-1). The percentage of GFP and scKoV DAg double-positive cells relative to DAPI-positive cells is shown. Cell numbers were quantified by averaging counts from five randomly selected microscopic fields per sample. Data are presented as the mean ± SD from three independent experiments (n = 3). Statistical significance was assessed by Tukey’s multiple-comparison test. ***p < 0.001, **p < 0.01, *p < 0.05. (**C**) Representative immunofluorescence images of cells after scKoV-WT transmission, acquired by fluorescence microscopy. Blue, DAPI; green, GFP; red, scKoV DAg; magenta, bornavirus nucleoprotein (N). Scale bar, 50 μm.

We also screened RNA samples previously used for avian bornavirus diagnosis for scKoV (*25–28*). Using RT-PCR and Sanger sequencing targeting a partial DAg region, we detected scKoV in 12 out of 18 CnBV-positive canary samples, whereas the only available CnBV-negative canary sample was scKoV-negative. Additionally, scKoV was not detected in the brain of an estrildid finch positive for estrildid finch bornavirus 1 (EsBV-1; *Orthobornavirus estrildidae*) (Fig. 4A and Table S5). Among parrot samples, scKoV was detected in samples from 5 of 77 tested birds. Of these five scKoV-positive birds, three were co-positive for PaBV-2, PaBV-5, or PaBV-4 and PaBV-7. No bornavirus had been detected in the remaining two birds under our screening conditions (Table S5).

Phylogenetic analysis including the obtained sequences showed that they clustered into the three known genotypes, with genotype 2 now further resolving into two sublineages, designated genotype 2a and genotype 2b (Fig. 4A). scKoV genotypes 1 and 2a were detected in passerines with CnBVs, whereas genotypes 3 and 2b were detected in parrots with or without diagnosed PaBV infections (Fig. 4A), except for genotype 1 detected in a *M. monachus* sample co-positive for PaBV-2 (SRR13480248). Together, these data further suggest that avian orthobornaviruses may be helper viruses for scKoV.

### 2.6. ORF2 promotes the spread of scKoV using avian bornaviruses as helper viruses

We next experimentally tested whether avian bornaviruses act as helper viruses for scKoV. We transfected pscKoV-WT into QT6 cells persistently infected with PaBV-2, PaBV-4, or munia bornavirus 1 (MuBV-1; *Orthobornavirus serini*) (producer cells). These cells were then co-cultured with puromycin-resistant GFP-expressing QT6 cells (recipient cells), and viral transmission was quantified after puromycin selection. Despite similar proportions of scKoV-positive producer cells (Fig. S13A), scKoV was detected in recipient cells only in the presence of avian bornaviruses (Figs. 4B and 4C). These results indicate that orthobornaviruses act as helper viruses for scKoV.

We next examined whether ORF2 is involved in the transmission of scKoV. We transfected pscKoV-ΔORF2 into QT6 cells persistently infected with PaBV-2, PaBV-4, or MuBV-1, and co-cultured these producer cells with the recipient QT6 cells (Fig. S13B). Although the co-infecting orthobornaviruses spread to nearly all recipient cells (Fig. S13C), scKoV-ΔORF2 transmitted less efficiently than scKoV-WT (Fig. 4B). Notably, scKoV transmission mediated by PaBV-2 or PaBV-4 was strongly dependent on ORF2, whereas MuBV-1-mediated transmission was less affected by ORF2 deletion (Fig. 4B). These results indicate that ORF2 promotes orthobornavirus-mediated scKoV transmission, although its importance differs among co-infecting viruses.

### 2.7. Ambisense gene expression and splicing in distinct lineages of KoVs

We investigated whether other KoVs also exhibit ambisense gene expression. Among the KoVs with genome sequences available in GenBank or reported by the Serratus project (Table S6) (*1*), five additional KoVs, including Indirana chiravasi deltavirus (ichiDV) and Desmodus rotundus deltavirus-B (DrDV-B), possess genomic-sense ORFs together with poly-A signals at plausible positions (Figs. 1A, 1B, and S14).

For ichiDV, the closest relative of scKoV, mapping analysis using a strand-specific RNA-seq dataset (SRR8954567) showed a clear ambisense transcription pattern similar to that of scKoV, although no clear evidence of splicing was detected (Fig. 1C). This suggests that ichiDV also uses an ambisense gene expression strategy. For other potential ambisense KoVs, mapping patterns did not show a clear bimodal distribution (Fig. S14). However, the corresponding RNA-seq libraries were prepared using rRNA depletion rather than poly-A enrichment. Thus, most viral reads were likely derived from genomic or antigenomic RNA rather than mRNA, making these RNA-seq datasets unsuitable for analyzing ambisense gene expression.

To further assess the transcriptional pattern of another potential ambisense KoV, we rescued DrDV-B in cell culture and performed strand-specific RNA-seq analysis (Fig. S15). Mapping analysis clearly showed an ambisense gene expression pattern and splicing in the 5’-UTR of the ORF2 transcript (Fig. 1C). These data indicate that DrDV-B also expresses ORF2.

The ichiDV ORF2 shows 36.6% amino acid sequence identity with scKoV ORF2a, and pairwise sequence alignment revealed that the N-terminal 80 residues are relatively well conserved (Fig. 1F). In contrast, DrDV-B ORF2 does not share significant amino acid sequence similarity with scKoV and ichiDV ORF2, nor does it show similarity in predicted secondary and tertiary structures (Figs. S9A and S9B).

## 3. Discussion

KoVs have been defined as negative-sense RNA viruses with small, highly self-complementary circular RNA genomes that impose strict constraints on genome organization and gene expression. Here, we showed that several KoVs, including scKoV, ichiDV, and DrDV-B, express the genomic-sense transcripts encoding a novel gene, ORF2, in addition to the canonical antigenomic DAg transcript (Fig. 1C). We also showed that scKoV expresses ORF2 during viral replication and that ORF2 is dispensable for viral replication but promotes helper virus-dependent transmission (Figs. 2A, 3A, and 4B). Together, this study provides novel insights into the genomic diversity of KoVs and suggests that KoVs can accommodate an additional gene despite strong constraints on their genomes.

Splicing may further expand the protein-coding capacity in some KoVs. Although KoVs transcribe in the nucleus, where host splicing machinery is available, direct evidence of splicing in KoVs has been lacking. Here, we showed that the ORF2 transcripts of scKoV and DrDV-B undergo splicing (Figs. 1C, 2C, and 2D). Interestingly, the biological significance of the splicing may differ between these two viruses. In scKoV, alternative splicing generates two ORF2 protein isoforms and thereby further expands the protein-coding capacity of the ORF2 gene. Indeed, the two ORF2 isoforms showed the distinct localization patterns in cells, suggesting functional differences between them (Fig. 3C). In contrast, based on the mapping pattern (Fig. 1C), splicing in DrDV-B occurs very efficiently in the 5’-UTR of ORF2, which may represent constitutive splicing. Thus, this splicing may help optimize ORF2 expression by removing regions required for maintaining genome self-complementarity but dispensable for protein function, thereby balancing structural constraints with efficient gene expression. Further studies will be required to identify further splicing patterns of KoVs.

Ambisense gene expression and splicing may have been independently acquired in distinct lineages of KoVs. These features are observed in phylogenetically distant KoV lineages represented by scKoV and DrDV-B, but not in other related KoVs, including HDV (Fig. 1A). Although repeated loss from a common ancestor cannot be excluded, independent acquisition of these traits provides a more parsimonious evolutionary explanation. This convergence suggests that ambisense transcription and alternative splicing constitute a feasible strategy for expanding coding capacity in KoVs while preserving their small, self-complementary circular genomes. Interestingly, this ambisense genome organization parallels that seen in other circular RNA replicons. Recently identified fungal ambiviruses encode two non-overlapping ORFs arranged in an ambisense orientation within a self-complementary region (*2*, *29*), resembling the genome organization of ambisense KoVs (Fig. 1B). Together, these similarities suggest convergent evolution of ambisense genome organization among distinct lineages of circular RNA replicons facing similar structural constraints. This also raises the possibility that additional ambisense circular RNA replicons remain to be discovered.

Our findings suggest that scKoV may regulate the balance between replication and helper-dependent transmission through a mechanism distinct from the A-to-I RNA editing strategy used by HDV. In HDV, A-to-I RNA editing at the DAg stop codon generates two DAg isoforms with distinct functions in replication and transmission (*8–11*, *13*). By contrast, we found no evidence that scKoV uses A-to-I RNA editing to generate alternative DAg isoforms. Instead, our data suggest that scKoV uses ambisense gene expression to express ORF2, a distinct factor that is dispensable for replication but promotes orthobornavirus-mediated transmission (Figs. 3A and 4B). Thus, scKoV may regulate the balance between replication and helper-dependent transmission through expression of an additional gene rather than through DAg isoform diversification. Whether this transmission-related role of ORF2 is conserved across other ambisense KoVs remains unknown. The molecular mechanism by which ORF2 promotes orthobornavirus-mediated transmission remains unclear and warrants further investigation.

We identified avian bornaviruses as helper viruses for scKoV. Most non-HDV KoVs lack identified helper viruses, with snake deltavirus as an exception that uses reptarenavirus (genus *Reptarenavirus*, family *Arenaviridae*) and/or hartmanivirus (genus *Hartmanivirus*, family *Arenaviridae*) as helper viruses (*19*, *30*). Using both public RNA-seq datasets and RT-PCR testing of diagnostic bird samples, we demonstrated co-infections of scKoV with CnBVs in passerines and PaBVs in psittacines (Fig. 4A). However, the non-representative availability of datasets and samples did not allow for the identifications of significant correlations. Nevertheless, co-infection with PaBV-2, PaBV-4, or MuBV-1 can support scKoV transmission in infected cell cultures (Figs. 4B and 4C), confirming their role as potential helper viruses. Interestingly, the extent of ORF2-dependent transmission varied depending on the helper viruses: ORF2 is less important for MuBV-1-mediated transmission than for PaBV-2- or PaBV-4-mediated transmission (Fig. 4B). This difference might reflect our use of scKoV genotype 1, which is predominantly associated with passerines, but this possibility will require future investigation. Notably, scKoV DAg showed altered subcellular localization in MuBV-1-infected cells (Figs. 4C and S13). HDV was reported to suppress the replication of HBV (*31*). Since both scKoV and MuBV-1 replicate in the nucleus, such interactions may influence viral replication including transcription, genome replication, and nuclear transport. Although the molecular basis of these interactions is unclear, our findings suggest that satellite–helper interactions in KoVs are more diverse and flexible than previously appreciated.

Co-infection with scKoV and avian bornaviruses also raises questions about the potential contribution of scKoV to disease. In humans, HDV co-infection exacerbates HBV-induced hepatitis and increases the risk of cirrhosis and hepatocellular carcinoma (*32*). Although the pathogenicity of scKoV remains unknown, it might also exacerbate diseases caused by co-infecting helper viruses. PaBVs are the causative agents of proventricular dilatation disease (PDD) in parrots (*33*, *34*), raising the possibility that scKoV could influence disease severity in co-infected birds. An even more intriguing case is that of CnBVs. Although CnBVs were reported to be associated with a PDD-like disease in canaries, experimental infection with CnBV isolates failed to reproduce the disease, suggesting that additional co-factors contribute to pathogenesis (*26*). In this context, scKoV might represent one such factor. Further studies will be needed to assess this possibility.

The mechanism underlying ambisense transcription remains unclear. Whereas scKoV, ichiDV, and DrDV-B show clear ambisense transcription, other KoVs, including Taeniopygia guttata DeV and Odocoileus virginianus DeV, appear to exhibit monosense transcription, expressing only the canonical antigenomic transcript encoding DAg (*15*). One possible explanation is that genomic-sense transcription depends on promoter activity on the antigenome, although no obvious differences in the predicted promoter regions were identified (Fig. 1D). The abundance of antigenomic templates may also influence the efficiency of ambisense transcription. Additionally, even when a genomic-sense transcript is produced, the absence of a downstream poly-A signal may reduce its stability. Indeed, most monosense KoVs lack a poly-A signal on the genomic strand. Defining the determinants of ambisense transcription will be important for understanding how KoVs diversify transcriptional strategies despite shared genomic constraints.

Taken together, our findings provide experimental evidence that circular RNA replicons can expand their coding capacity through ambisense gene expression despite stringent structural constraints on their genome architecture. Additionally, our results show that such ambisense gene expression can generate novel gene products that contribute to helper virus-dependent transmission in circular RNA satellites. These findings broaden our understanding of the evolutionary and functional diversity of circular RNA replicons and suggest that additional gene expression strategies, as well as previously unrecognized satellite-helper virus relationships, remain to be discovered.

## 4. Materials and Methods

### 4.1. Detection of scKoV-derived reads from public RNA-seq data

RNA-seq datasets from birds excluding domestic chickens (*Gallus gallus* [Linnaeus, 1758]) were downloaded from the NCBI Sequence Read Archive (SRA) (*35*) (accessed on Jan 20, 2024) and mapped to the partial scKoV genome (BR001664.1) using Magic-BLAST v1.7.1 (*36*) with the options “-splice T-word_size 16 -lcase_masking”. The mapped read numbers were counted using SAMtools v1.19 (*37*). The mapping results were manually inspected, and reads aligned to less than 25 bases of the reference sequence were excluded as non-specific.

### 4.2. Identification of complete scKoV genome sequences

The scKoV-positive RNA-seq datasets were downloaded and preprocessed using fastp v0.23.4 (*38*) with the options “-l 35 -x -y”. The preprocessed reads were mapped to the corresponding host genomes (Table S7) using HISAT2 v2.2.1 (*39*) with the default settings. The unmapped reads were extracted using SAMtools v1.19 and subjected to *de novo* assembly using Trinity v2.15.1 (*40*) with the default settings. The scKoV contigs were identified through a two-step sequence similarity search: BLASTn against the partial scKoV genome and BLASTx against the clustered nr database (*35*), using BLAST+ v2.15.0 (*41*).

Self-dotplot analysis of the linear scKoV contigs was performed using the YASS program with the default settings (*42*) (Fig. S1). Contigs with identical sequences at both termini were manually circularized using Geneious Prime v2024.0.5 (https://www.geneious.com). The circularity of scKoV contigs was further verified by mapping short reads to the circularized scKoV contigs using Geneious and manually checking the mapped reads across the circularized boundaries.

For scKoV-positive RNA-seq datasets in which circular contigs could not be obtained by *de novo* assembly, reads were mapped to the circular scKoV contigs using Geneious, and circular consensus sequences were extracted. The depth of mapped reads was counted for all contigs using SAMtools v1.19 (*37*). Each position was confirmed to be covered with at least three reads.

### 4.3. Identification of complete genome sequences of known KoVs

For KoVs identified in the Serratus project for which complete genome sequences were publicly unavailable (*1*), complete genome sequences were independently reconstructed. SRA accession numbers of RNA-seq datasets containing reads alignable to deltavirus DAg sequences were obtained from the Serratus Explorer (https://serratus.io/, accessed on Apr 9, 2025). The RNA-seq datasets were assembled, and KoV sequences were identified and characterized as described in section 4.2. Information on the identified complete KoV sequences is shown in Table S6. Unexpectedly, a KoV sequence obtained from an RNA-seq datasets of a mixed Lepidoptera sample (SRR3400941) was identical to that of a previously reported KoV from the Chusan Island toad (*20*). However, metadata and read-mapping analysis showed that the RNA-seq data originated from insects of the order Lepidoptera (see Supporting text 2).

### 4.4. Characterization of KoV genomes

The self-complementarities of KoV genomes were analyzed using the Mfold web server (*43*) with the default settings. Genomic and antigenomic ribozyme structures were initially inferred using the IPknot web server (*44*), with the visualized output from IPknot serving as a guide for manual ribozyme structure drawing. Poly-A signals (AAUAAA) were manually identified. ORFs were detected using the ‘Find ORF’ function in Geneious, with a minimal threshold of 200 nucleotides. ORFs that overlap with the terminal rod-like structure of the viral genome, where the putative transcriptional promoter is located, poly-A signals, or the predicted ribozymes were excluded.

### 4.5. Short read mapping to analyze transcriptional patterns

The short reads from KoV-positive RNA-seq data were preprocessed as described above (see the section 4.2) and mapped to the corresponding KoV genomes using HISAT2 v2.2.1 (*39*) with the options “-k 2 --max-intronlen 1000 --rna-strandness RF”. To accurately evaluate reads spanning the circularized genome boundaries, 2× tandemly repeated genomes were used as the references. The “-k 2” option in HISAT2 was selected to ensure precise mapping to the 2× genomes. For strand-specific RNA-seq data, the “--rna-strandness RF” option was used to retain strand information. The mapping patterns were visualized using Geneious.

### 4.6. Phylogenetic analysis

Phylogenetic relationship among all known KoVs was inferred as follows. The amino acid sequences of KoV DAg were aligned by MAFFT v7.490 with the E-INS-i algorithm (*45*). The phylogenetic tree was reconstructed using the Bayesian Markov chain Monte Carlo (MCMC) method in MrBayes v3.2.7a (*46*) with two independent runs, four chains per run, and the JTT model of substitution. The analysis was run for 5 million iterations, with samples taken every 5,000 steps. The average standard deviation of split frequencies was 0.010.

For the full-length scKoV genomes and partial DAg sequences, nucleotide sequences were aligned using MAFFT v7.490 with the E-INS-i algorithm (*45*). Pairwise sequence identities were calculated from the alignments using Geneious. Maximum-likelihood trees were inferred in RAxML v1.1.0 (*47*) using substitution models selected by the minimum BIC (TVM+I+G4 model for full-length genome and HKY+I for partial DAg sequences). The reliability of the trees was assessed by 1,000 bootstrap resampling with the transfer bootstrap expectation method (*48*). The trees were visualized and annotated in Interactive Tree Of Life (iTOL) v6.9.1 (*49*) or MEGA 12.1 (*50*). The alignments used for the phylogenetic analyses are provided in the Supplementary Materials.

### 4.7. Detection of RNA-editing at the DAg stop codon

Sequence variations at the DAg stop codon were identified from the mapping data described in Section 4.5 using the ‘Find SNP/Variants’ function in Geneious.

### 4.8. In silico characterization of ORF2 proteins

The amino acid sequences of both ORF2a and ORF2b were analyzed using BLASTp, with the PSI-BLAST and the DELTA-BLAST algorithms and an *E*-value threshold of 0.05. Theoretical molecular weight and isoelectric point were calculated using the ProtParam web server (https://web.expasy.org/protparam/). Amino acid sequence alignments, colored according to hydrophobicity and polarity, were generated using Geneious. Membrane topology of ORF2 proteins was inferred using DeepTMHMM ver. 1.0 (*51*). Nuclear localization signals were inferred using the cNLS mapper web server with a cut-off value of 3.0 (*52*). Intrinsically disordered regions were predicted using AIUPred web server (*53*). Secondary structures of ORF2 proteins were predicted using JPred v4 (*54*). Protein structure predictions were performed by the AlphaFold3 web server (*55*).

### 4.9. Cell culture

QT6 (quail fibroblast) cells and LMH (chicken hepatocellular carcinoma) cells were maintained in Dulbecco’s modified Eagle’s medium (DMEM)/Ham’s F-12 (11581-15; Nacalai Tesque, Kyoto, Japan) supplemented with 5% fetal bovine serum (FBS; Sigma-Aldrich, St. Louis, MO, USA) and 10% FBS, respectively, and incubated at 37°C in a humidified 5% CO_2_ atmosphere. DF-1 (chicken fibroblast) cells were cultured in low-glucose DMEM (08456-36; Nacalai Tesque) supplemented with 10% FBS and incubated at 39°C under the same atmospheric conditions.

### 4.10. Antibody production

The ORFs encoding DAg and ORF2 of scKoV (BR002461) were cloned into pET-15b expression vector. *Escherichia coli* Rosetta2(DE3)pLysS cells were transformed with the resulting plasmids and cultured in LB medium supplemented with 100 µg/mL ampicillin and 34 µg/mL chloramphenicol at 37°C. When the cultures reached an OD₆₀₀ of 0.4–0.6, protein expression was induced with 0.4 mM IPTG and the cultures were incubated at 37°C for 3 h. The soluble fraction obtained after sonication was purified using TALON metal affinity resin (Takara Bio, Shiga, Japan).

Immunization of mice with the purified DAg protein and antiserum production were performed by Cosmo Bio (Tokyo, Japan). Immunization of rabbits with the purified DAg or ORF2 proteins and antiserum production were carried out by Eurofins Genomics (Tokyo, Japan). The rabbit antisera were subjected to ammonium sulfate precipitation and dialyzed against PBS.

### 4.11. Indirect immunofluorescence assay (IFA)

Cells were fixed with 4% paraformaldehyde phosphate buffer solution (FUJIFILM Wako, Osaka, Japan) and permeabilized with 0.4% Triton X-100 (Nacalai Tesque) in PBS. After blocking with PBS containing 0.1% Tween 20 and 1% bovine serum albumin (FUJIFILM Wako), the cells were incubated with primary antibodies against scKoV DAg, ORF2, and/or Borna disease virus 1 N protein (HB01). Then, the cells were incubated with Alexa Fluor 488-, 555-, and/or 680-conjugated secondary antibodies (Thermo Fisher Scientific, Waltham, MA, USA) and DAPI (FUJIFILM Wako). Primary and secondary antibodies, including their sources, catalog numbers, dilutions, and incubation conditions, are listed in Table S8. Fluorescence signals were visualized using an EVOS M7000 fluorescence microscope (Thermo Fisher Scientific) or a FV3000 confocal microscope (Olympus, Tokyo, Japan).

### 4.12. Western blotting

Cells were lysed in SDS sample buffer [125 mM Tris-HCl (pH 6.8), 4% SDS, 10% sucrose, 10% dithiothreitol (DTT), and 0.01% bromophenol blue] and the lysates were subjected to SDS-PAGE, followed by transfer to polyvinylidene difluoride (PVDF) membranes (Merck, Darmstadt, Germany). The membranes were blocked with 5% skim milk (FUJIFILM Wako) and incubated with primary antibodies against scKoV DAg and ORF2. After washing, the membranes were probed with horseradish peroxidase (HRP)-conjugated secondary antibodies (Jackson ImmunoResearch Laboratories, West Grove, PA, USA). Primary and secondary antibodies, including their sources, catalog numbers, dilutions, and incubation conditions, are listed in Table S8. Signals were detected using Chemi-Lumi One L (Nacalai Tesque).

### 4.13. Rescue of scKoV

A plasmid encoding a 1.2× negative-strand genome of scKoV genotype 1 (BR002461), spanning from 40 bp upstream of the putative genomic ribozyme to 40 bp downstream of the putative antigenomic ribozyme, was synthesized in pTwist-CMV vector (pTwist-CMV-1.2×scKoV) by Twist Bioscience (San Francisco, CA, USA). Plasmids for DAg and ORF2-deficient mutants (pTwist-CMV-1.2×scKoV-ΔDAg and pTwist-CMV-1.2×scKoV-ΔORF2, respectively) were constructed by inverse PCR, in which the start codons (ATG) of DAg and ORF2 were individually mutated to GTG.

To rescue scKoV and its mutants, the constructed plasmids were transfected into cells using Avalanche-Everyday Transfection Reagent (APRO Science, Tokushima, Japan). The transfected cells were passaged and maintained until used for IFA and western blotting. The intended mutations were confirmed by RT-PCR and Sanger sequencing as described in Section 4.14.

### 4.14. Endpoint RT-PCR

Total RNA was extracted from cells using NucleoSpin RNA Plus kit (Macherey-Nagel, Düren, Germany). Residual DNA was removed with rDNase (Macherey-Nagel). Reverse transcription was performed using Verso cDNA Synthesis Kit (Thermo Fisher Scientific) with Anchored Oligo dT primer to detect alternative splicing or using ReverTra Ace qPCR RT Master Mix (TOYOBO, Osaka, Japan) to verify mutation site in scKoV mutants. PCR was performed with Phusion Green Hot Start II High-Fidelity Master Mix (Thermo Fisher Scientific) with the primers listed in Table S9. The PCR products were purified and subjected to Sanger sequencing by Eurofins Genomics (Tokyo, Japan).

### 4.15. Subcellular localization of scKoV ORF2 isoforms

pTwist-CMV plasmids encoding either ORF2a, ORF2b, or ORF2 WT were generated by seamless cloning and inverse PCR (Fig. S11). QT6 cells were transfected with the resulting plasmids, and ORF2 expression was analyzed by western blotting and IFA.

### 4.16. Detection of co-existing viruses

To identify co-existing viruses in scKoV-positive RNA-seq datasets, the assembled contigs (section 4.2) were clustered using CD-HIT v4.8.1 (*56*) with the 95 % threshold. The clustered contigs of 500 nucleotides or longer were extracted using SeqKit v2.4.0 (*57*), which were subjected to a two-step sequence similarity search. The first search was conducted by MMseqs2 (version c48da9d781b81804727b5cccfed7f97cfcc20c9d) (*58*) with the default settings against a custom database consisting of viral protein sequences with a threshold *E*-value of 0.001. The custom database was created as follows. Viral protein sequences were downloaded from the NCBI nr database (*35*) on June 22, 2023, which was clustered by using CD-HIT (*56*) with a threshold of 0.9. Contigs alignable to viral sequences other than retroviruses and environmental viruses were extracted. The second BLASTx search was then performed using the extracted contigs as queries against the NCBI clustered nr database (*35*) with the options “-e-value 1e-4 -word_size 2 -max_target_seqs 10” on Aug 26, 2024. The contigs with the best hit to viruses were extracted and manually analyzed. Contigs with the best hit to bornaviruses (family *Bornaviridae*) were further subjected to a BLASTn search against a database consisting of CnBVs and PaBVs to accurately identify the species/genotypes.

To quantify avian bornavirus-derived reads in scKoV-positive RNA-seq datasets, preprocessed reads were mapped using HISAT2 (*39*) with the option “-k 1” against CnBV-1 (NC_030690.1), CnBV-3 (NC_024296.1), and newly obtained PaBV-2 contig assembled from SRR13480248 (Supplementary Materials) because the genome sequence of coexisting PaBV-2 is slightly different (91.97 % nt identity; vs FJ620690) from the those deposited in INSDC.

### 4.17. scKoV screening using diagnostic bird samples

RNA extracted from diagnostic samples originating from passerine and psittacine birds was tested for scKoV by conventional RT-PCR. The samples had been previously tested for CnBVs and PaBVs (Table S5) (*25–28*).

RT-PCR was performed using the SuperScript III One-Step RT-PCR System with Platinum Taq DNA Polymerase (Thermo Fisher Scientific) and the primers scKoV-1F and scKoV-2R (Table S9). RNA and primers were first incubated at 70°C for 5 min and immediately chilled at 4°C for 1 min, after which the remaining reaction components were added. The PCR reactions were performed under the following conditions: 60°C for 30 min; 94°C for 2 min; 40 cycles of 94°C for 15 sec, 55°C for 30 sec, and 68°C for 30 sec; and a final extension at 68°C for 3 min. PCR products were analyzed by agarose gel electrophoresis, and bands of the expected size were purified and subjected to Sanger-sequencing at Eurofins Genomics (Ebersberg, Germany). The resulting primer-trimmed sequences (length 197 bp) were submitted to GenBank under accession numbers LC923643 to LC923659.

### 4.18. Transmission assay of scKoV in the presence or absence of avian bornaviruses

Uninfected QT6 cells or QT6 cells persistently infected with PaBV-2, PaBV-4, or MuBV-1 were transfected with pTwist-CMV-1.2×scKoV or pTwist-CMV-1.2×scKoV-ΔORF2 to establish producer cells infected with scKoV alone or co-infected with scKoV and each avian bornavirus. The resulting producer cells were mixed at a 1:1 ratio with uninfected puromycin-resistant, GFP-expressing QT6 cells and co-cultured for 14 days. To selectively retain GFP-positive recipient cells, puromycin was added to the culture medium at a final concentration of 10 µg/mL for 3 days. After selection, the cells were subjected to indirect IFA to assess the expression of scKoV and bornavirus proteins.

## Acknowledgements

We would like to thank Kathrin Steffen for excellent technical assistance. The antibody against Borna disease virus 1 N protein (HB01) was kindly provided by Dr. Keizo Tomonaga (Institute for Life and Medical Sciences, Kyoto University, Kyoto, Japan).

The super-computing resources were provided by Human Genome Center, the Institute of Medical Science, the University of Tokyo.

## Author contributions

Conceptualization: M.K., M.H.; Methodology: M.K., M.I., M.H.; Formal analysis: M.K.; Investigation: M.K., S.I., A.S.; Resources: Y.S.-O., D.R., M.H.; Data curation: M.K.; Writing – original draft: M.K.; Writing – review & editing: M.K., S.I., A.S., M.I., Y.S.-O., D.R., M.H.; Visualization: M.K.; Supervision: M.K., M.H.; Project administration: M.K., M.H.; Funding acquisition: M.K., M.I., M.H.

## Data and materials availability

The sequence data have been deposited in GenBank under accession numbers XXXX–XXXX. The raw RNA-seq data are available in the DDBJ Sequence Read Archive under accession numbers DRAxxxxxx.

