## Supporting Text for "Ambisense transcription uncovers hidden coding capacity in compact circular RNA satellite viruses"

***1. No functional domains were identified in ORF2 proteins***

To investigate the potential function of ORF2 proteins, we performed *in silico* analyses using sequence similarity searches, structural predictions, and functional domain searches. Neither ORF2a nor ORF2b showed detectable similarity to known protein families or conserved domains. Secondary- and tertiary-structure prediction suggested an N-terminal α-helix and an unstructured C-terminus, but no reliable full-length structure was predicted (Fig. S9A and S9B). An intrinsically disordered region was detected only in the C-terminal third of ORF2b (Fig. S9C). Additionally, no obvious hydrophobic or polar regions were identified (Figs. S9D and S9E). Predictive analyses of nuclear localization signals and transmembrane domains also yielded no significant features (Fig. S9F). These results suggest that the ORF2 gene encodes proteins of previously uncharacterized evolutionary origin.

***2. Chusan Island toad virus 1 was detected from mix sample of Lepidoptera spp.***

A KoV sequence obtained from a mixed Lepidoptera RNA-seq sample (SRR3400941) was found to be identical to a previously reported Chusan Island toad virus 1 (*1*). To determine whether this RNA-seq dataset originated from frogs/toads or moths/butterflies, we mapped reads to DNA barcodes sequence for species classification. The 5′ region of mitochondrial cytochrome c oxidase subunit I (COI-5P) barcode sequences of the orders Anura and Lepidoptera were downloaded from Bold Systems v4 (https://v4.boldsystems.org/, accessed on October 4, 2025) and clustered at 98% sequence identity using CD-HIT v4.8.1. The RNA-seq dataset (SRR3400941) was mapped to the clustered COI-5P barcodes using Magic-BLAST v1.7.2, and the mapped reads were counted with SAMtools v1.16.1. A total of 26,955 reads were mapped to Lepidoptera, whereas only 13 reads were mapped to Anura (Table S10). This finding suggests that SRR3400941 was derived from Lepidoptera rather than Anura, raising the possibility that Chusan Island toad virus 1 may also originate from moths or butterflies.
