## Supplementary Figures for "Ambisense transcription uncovers hidden coding capacity in compact circular RNA satellite viruses"

### Fig. S1

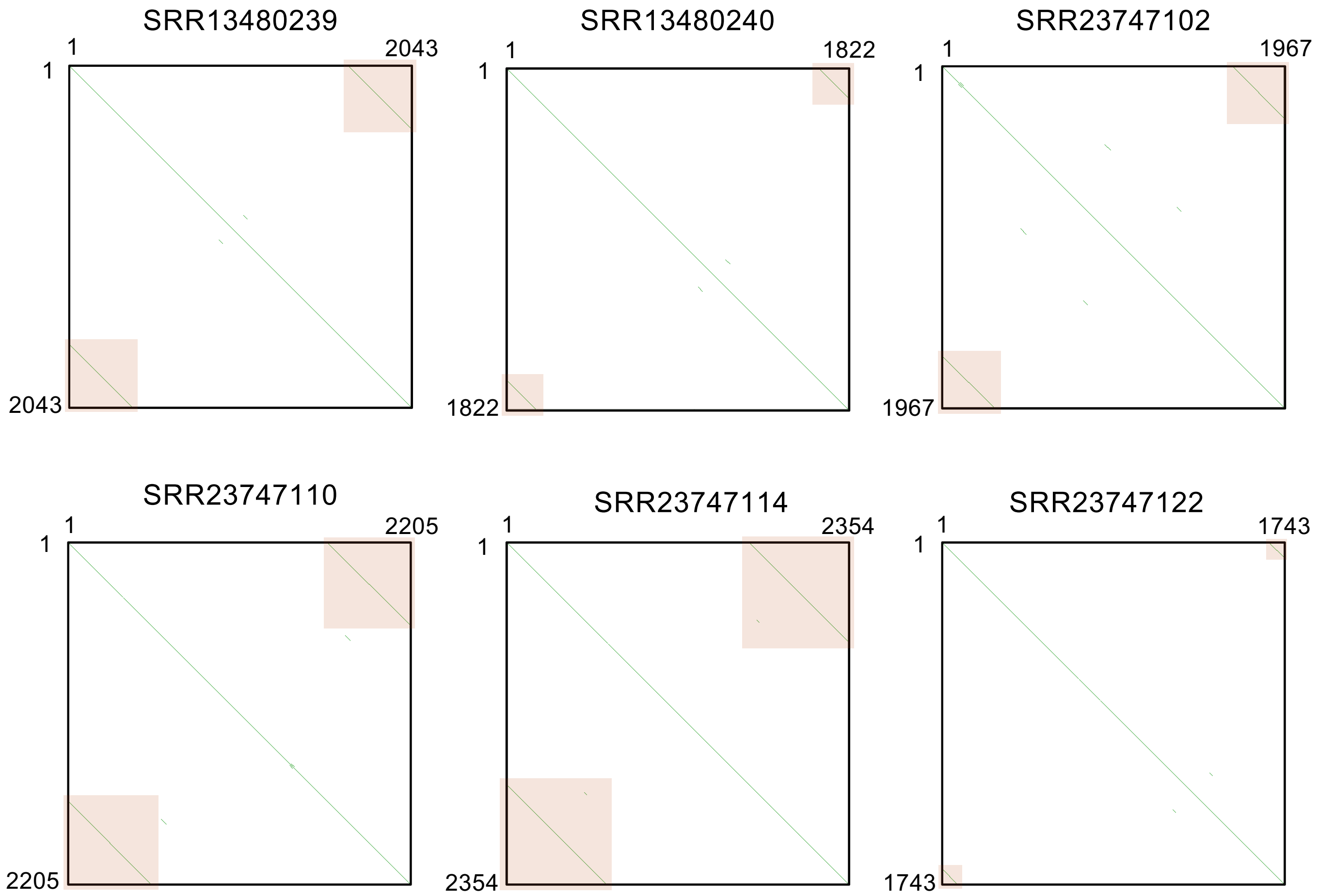


**Supplementary Figure 1. Self-dotplot analyses of scKoV contigs.**
Each scKoV contig was analyzed using the YASS program (Noé et al. 2005). Numbers indicated nucleotide positions of the contigs. Green lines indicate the sequences aligned in the forward direction. Contig ends with identical sequences are highlighted by orange boxes, indicating that the assembled contigs derive from complete circular genomes.

### Fig. S2

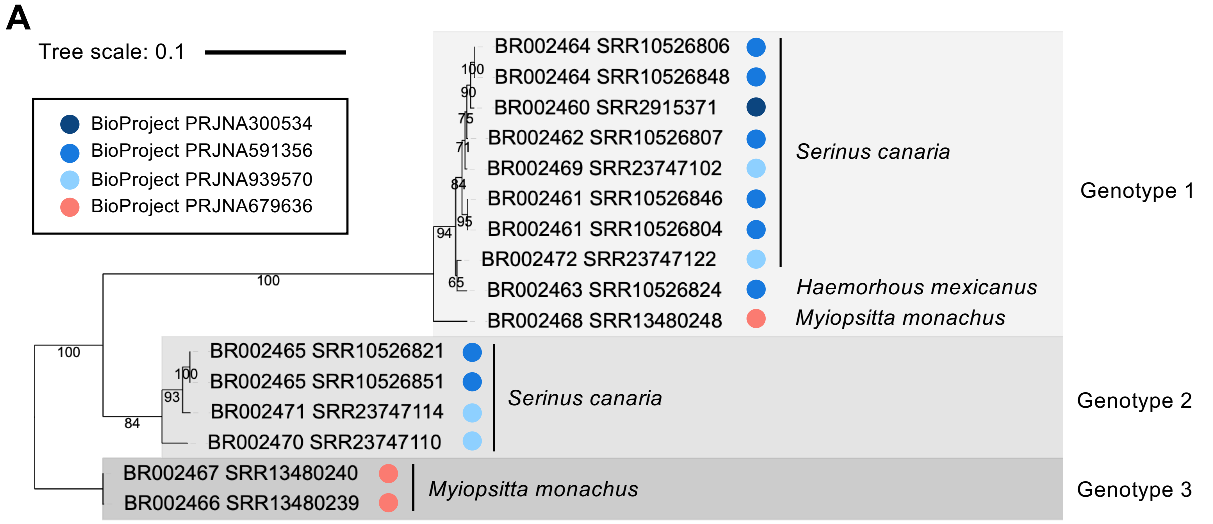


**B**

DAg (aa) (upper right); Genome (nt) (lower left); Identity (%): High – Low

| | | 1 | 2 | 3 | 4 | 5 | 6 | 7 | 8 | 9 | 10 | 11 | 12 | 13 | 14 | 15 | 16 |
|---|---|---|---|---|---|---|---|---|---|---|---|---|---|---|---|---|---|
| 1 | BR002460 SRR2915371 *S. canaria* | | 100.0 | 100.0 | 100.0 | 99.5 | 100.0 | 100.0 | 99.5 | 99.5 | 97.6 | 93.2 | 93.2 | 92.7 | 93.7 | 91.7 | 92.2 |
| 2 | BR002464 SRR10526806 *S. canaria* | 99.5 | | 100.0 | 100.0 | 99.5 | 100.0 | 100.0 | 99.5 | 99.5 | 97.6 | 93.2 | 93.2 | 92.7 | 93.7 | 91.7 | 92.2 |
| 3 | BR002464 SRR10526848 *S. canaria* | 99.5 | 100.0 | | 100.0 | 99.5 | 100.0 | 100.0 | 99.5 | 99.5 | 97.6 | 93.2 | 93.2 | 92.7 | 93.7 | 91.7 | 92.2 |
| 4 | BR002462 SRR10526807 *S. canaria* | 99.5 | 99.4 | 99.4 | | 99.5 | 100.0 | 100.0 | 99.5 | 99.5 | 97.6 | 93.2 | 93.2 | 92.7 | 93.7 | 91.7 | 92.2 |
| 5 | BR002469 SRR23747102 *S. canaria* | 99.3 | 99.2 | 99.2 | 99.7 | | 99.5 | 99.5 | 99.0 | 99.0 | 97.1 | 92.7 | 92.7 | 92.7 | 93.2 | 91.3 | 91.7 |
| 6 | BR002461 SRR10526804 *S. canaria* | 99.2 | 99.2 | 99.2 | 99.2 | 99.3 | | 100.0 | 99.5 | 99.5 | 97.6 | 93.2 | 93.2 | 92.7 | 93.7 | 91.7 | 92.2 |
| 7 | BR002461 SRR10526846 *S. canaria* | 99.2 | 99.2 | 99.2 | 99.2 | 99.3 | 100.0 | | 99.5 | 99.5 | 97.6 | 93.2 | 93.2 | 92.7 | 93.7 | 91.7 | 92.2 |
| 8 | BR002463 SRR10526824 *H. mexicanus* | 98.3 | 98.3 | 98.3 | 98.6 | 98.7 | 98.4 | 98.4 | | 100.0 | 98.1 | 92.7 | 92.7 | 92.2 | 93.2 | 91.3 | 91.7 |
| 9 | BR002472 SRR23747122 *S. canaria* | 98.8 | 98.5 | 98.5 | 98.9 | 98.9 | 98.8 | 98.8 | 99.0 | | 98.1 | 92.7 | 92.7 | 92.2 | 93.2 | 91.3 | 91.7 |
| 10 | BR002468 SRR13480248 *M. monachus* | 95.6 | 95.3 | 95.3 | 95.6 | 95.6 | 95.9 | 95.9 | 95.6 | 95.9 | | 92.2 | 92.2 | 91.7 | 92.7 | 90.8 | 91.3 |
| 11 | BR002465 SRR10526821 *S. canaria* | 83.0 | 82.8 | 82.8 | 83.1 | 83.1 | 83.3 | 83.3 | 83.0 | 83.3 | 83.0 | | 100.0 | 99.5 | 99.5 | 93.7 | 94.2 |
| 12 | BR002465 SRR10526851 *S. canaria* | 83.0 | 82.8 | 82.8 | 83.1 | 83.1 | 83.3 | 83.3 | 83.0 | 83.3 | 83.0 | 100.0 | | 99.5 | 99.5 | 93.7 | 94.2 |
| 13 | BR002471 SRR23747114 *S. canaria* | 83.3 | 83.0 | 83.0 | 83.1 | 83.1 | 83.4 | 83.4 | 83.0 | 83.4 | 83.0 | 99.0 | 99.0 | | 99.0 | 93.2 | 93.7 |
| 14 | BR002470 SRR23747110 *S. canaria* | 83.4 | 83.3 | 83.3 | 83.5 | 83.6 | 83.8 | 83.8 | 83.1 | 83.6 | 83.1 | 96.5 | 96.5 | 96.4 | | 94.2 | 94.7 |
| 15 | BR002466 SRR13480239 *M. monachus* | 81.6 | 81.7 | 81.7 | 81.9 | 82.0 | 82.1 | 82.1 | 81.7 | 81.8 | 82.1 | 88.5 | 88.5 | 88.6 | 88.5 | | 99.5 |
| 16 | BR002467 SRR13480240 *M. monachus* | 81.7 | 81.7 | 81.7 | 82.0 | 82.0 | 82.2 | 82.2 | 81.8 | 81.9 | 82.1 | 88.5 | 88.5 | 88.6 | 88.5 | 99.9 | |

**Supplementary Figure 2. Phylogenetic analysis of scKoV variants.** (A) The phylogenetic tree was inferred by the maximum likelihood method using nucleotide sequences of complete scKoV genomes. Bootstrap values are indicated at each branch. The BioProject from which each scKoV sequence was obtained is indicated by a colored circle. The scale bar indicates the number of nucleotide substitutions per site. (B) Heat map of pairwise identities of scKoV variants based on nucleotide sequences of complete genomes (lower left) and amino acid sequences of DAg (upper right).

Fig. S3

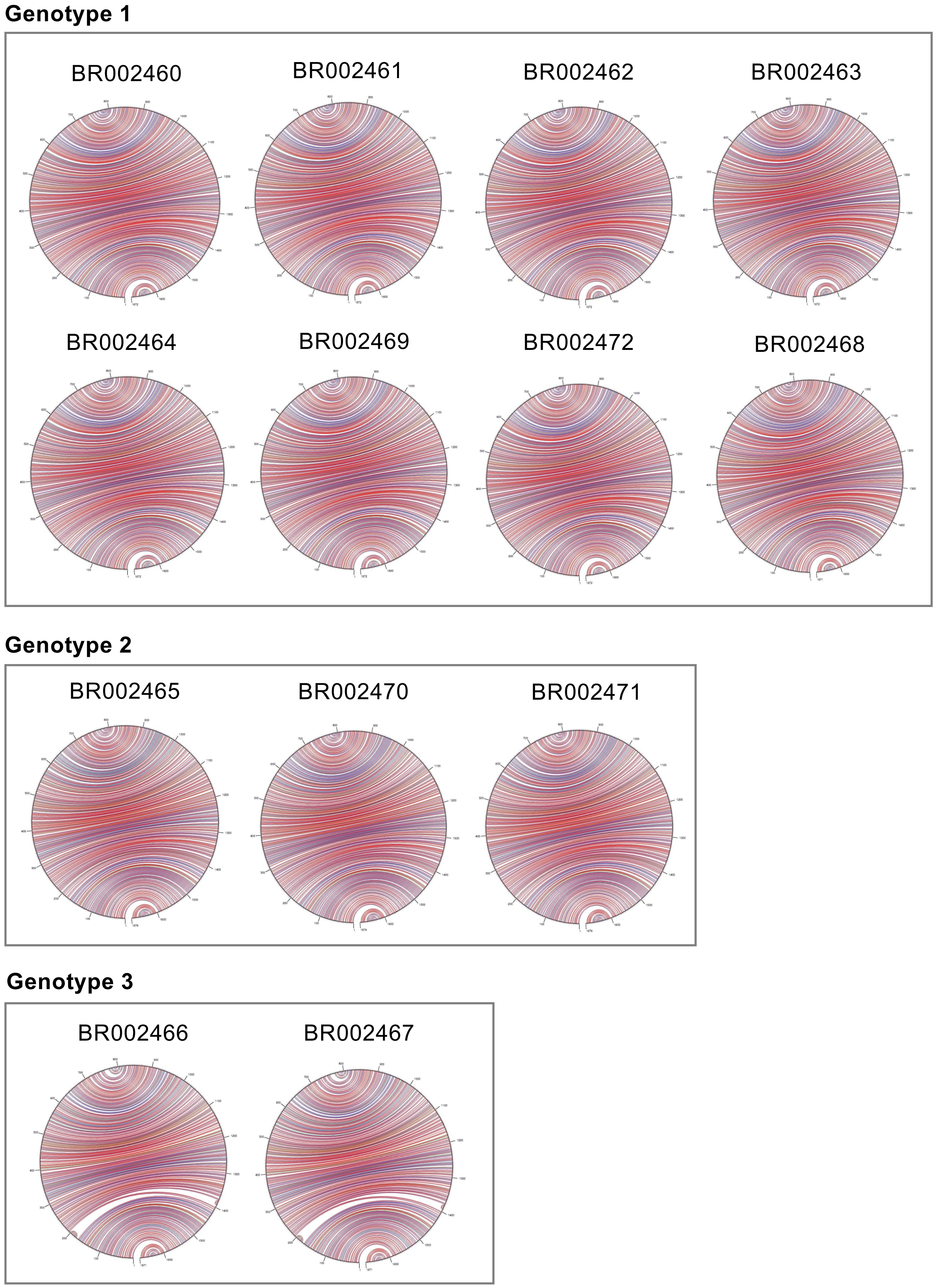


**Supplementary Figure 3. Self-complementarity analyses of the scKoV genomes.**
RNA secondary structures of the complete scKoV genomes were predicted using Mfold web server (Zuker et al. 2003). Red, blue, and green arcs indicate G-C, A-U, and G-U pairs, respectively. Information on scKoV genotypes is shown in Fig. 4A and Extended Data Fig. 1.

Fig. S4

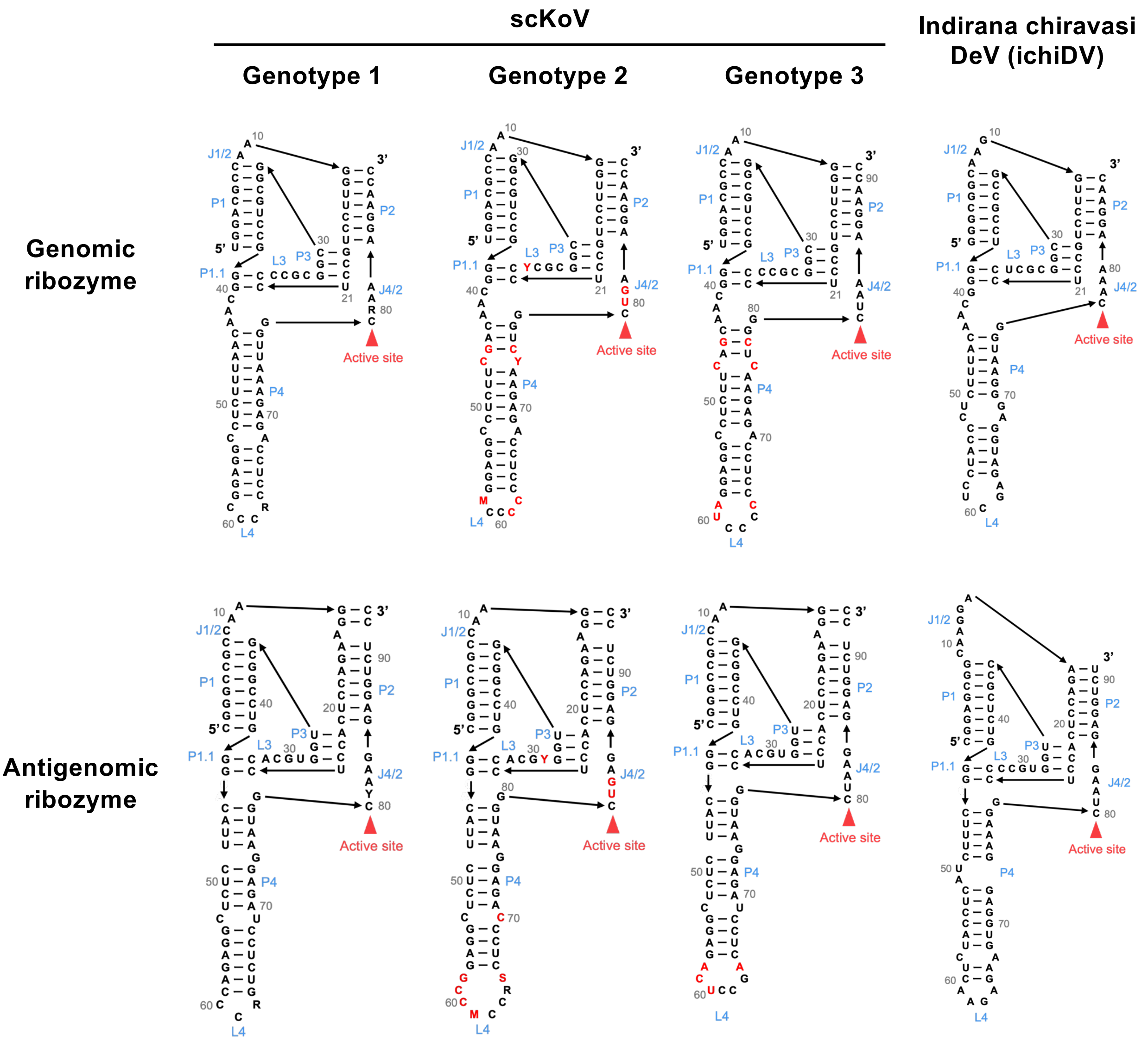


**Supplementary Figure 4. Predicted structure of genomic and antigenomic ribozymes.** Predicted genomic and antigenomic ribozyme structures of scKoV genotypes 1, 2, and 3 and Indirana chiravasi DeV (ichiDV). Gray numbers indicate nucleotide positions of each predicted ribozyme. Consecutive bases are connected by arrows from 5' to 3'. Sequence variations within each genotype are indicated by mixed bases. The name of secondary structural elements are shown in blue. Catalytic sites are indicated by red triangles. Sequence differences from genotype 1 are highlighted in red.

Fig. S5

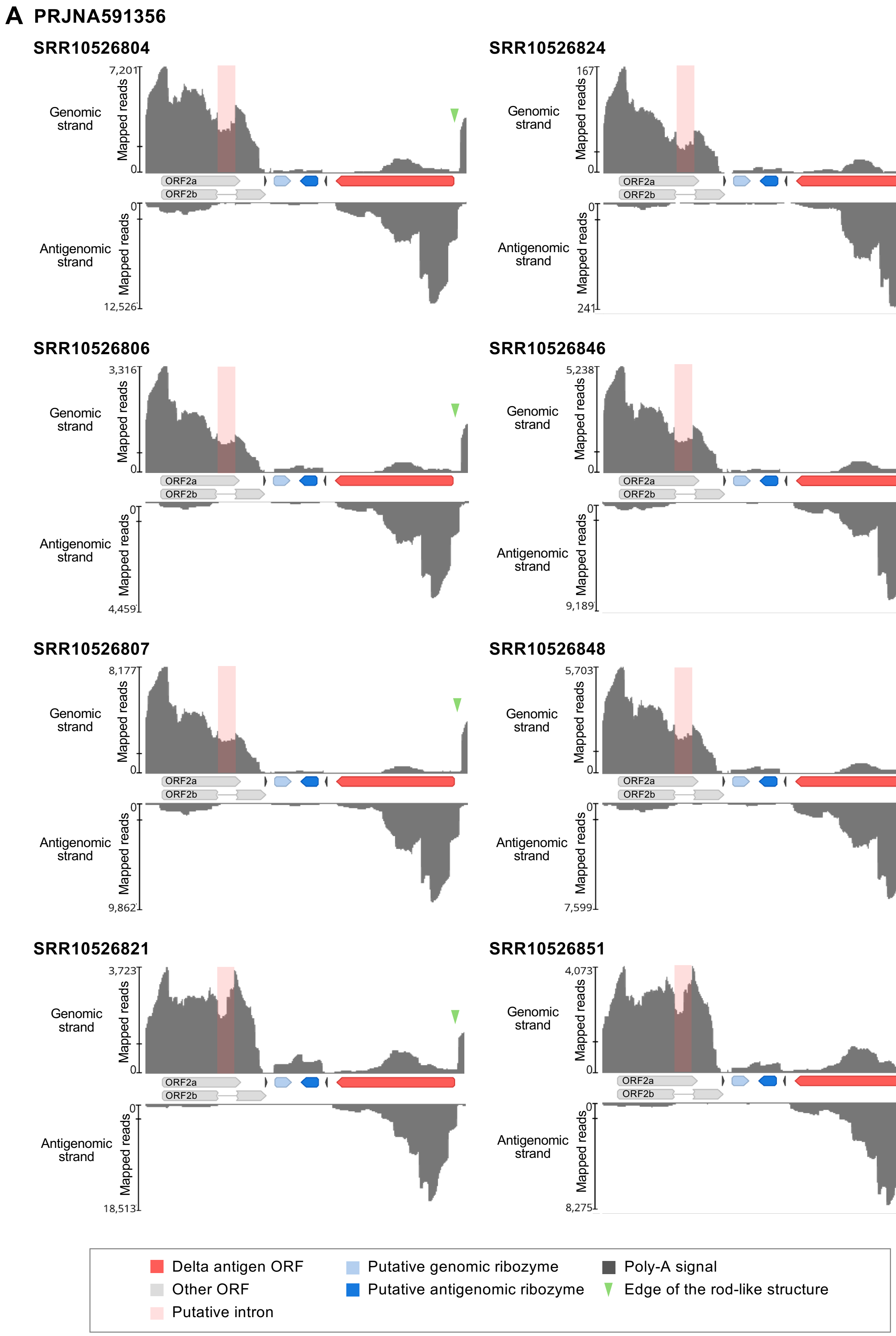
A PRJNA591356
SRR10526804
7,201
0
12,526
Genomic strand
Antigenomic strand
Mapped reads
ORF2a
ORF2b
SRR10526824
167
241
SRR10526806
3,316
4,459
SRR10526846
5,238
9,189
SRR10526807
8,177
9,862
SRR10526848
5,703
7,599
SRR10526821
3,723
18,513
SRR10526851
4,073
8,275
Delta antigen ORF
Other ORF
Putative intron
Putative genomic ribozyme
Putative antigenomic ribozyme
Poly-A signal
Edge of the rod-like structure

Fig. S5

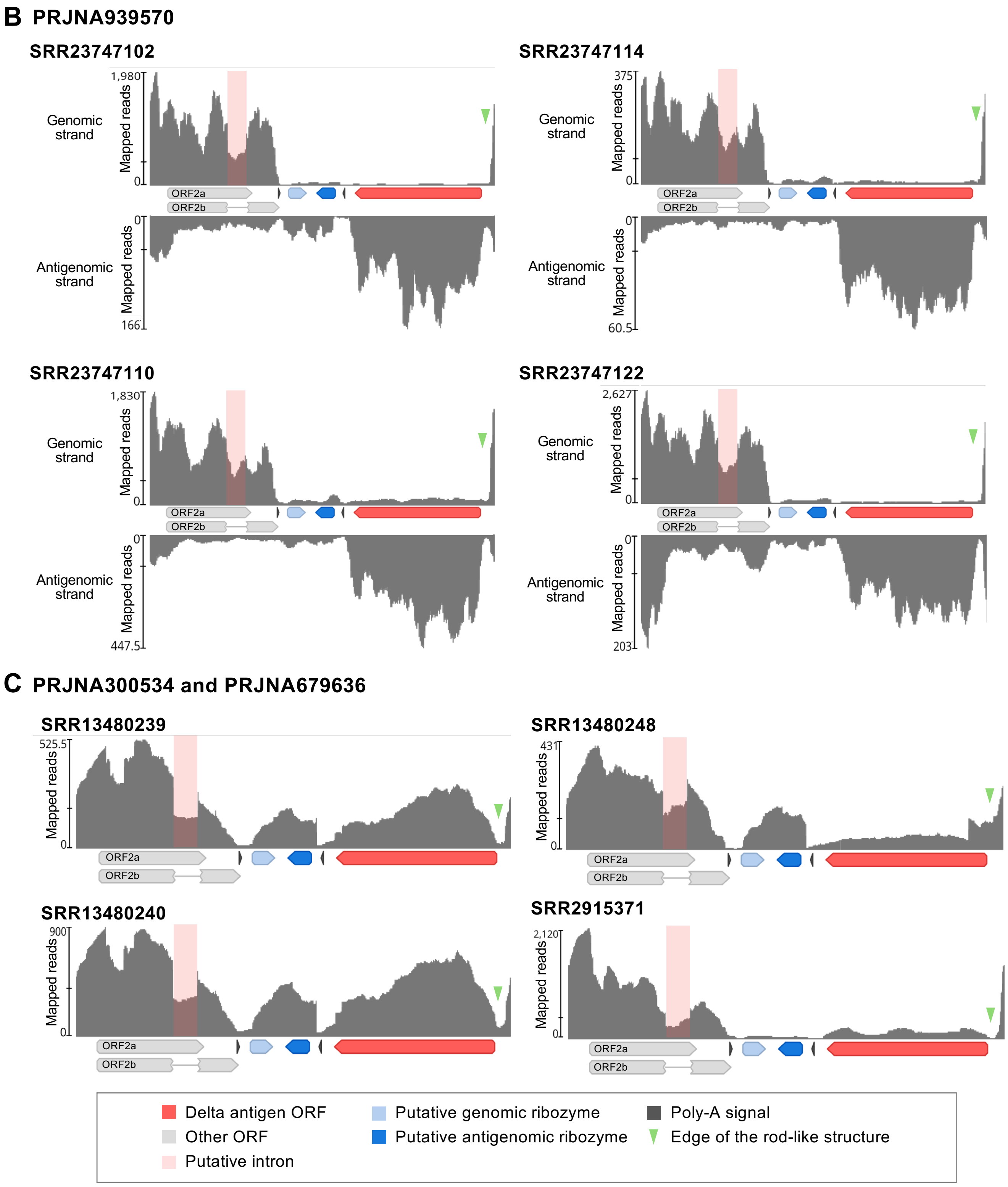


**Supplementary Figure 5. Transcriptional pattern of scKoV.**

(**A and B**) Strand-specific mapping coverage of original short reads of scKoV complete genome sequence in BioProjects PRJNA591356 (A) and PRJNA939570 (B). (**C**) Non-strand-specific mapping coverage of original short reads of scKoV complete genome sequence. Colored arrows indicate annotations (ORFs, ribozymes, and poly-A signals). Light pink boxes indicate putative introns. Green triangle indicates edge of the rod-like structure of circular scKoV genome.

Fig. S6

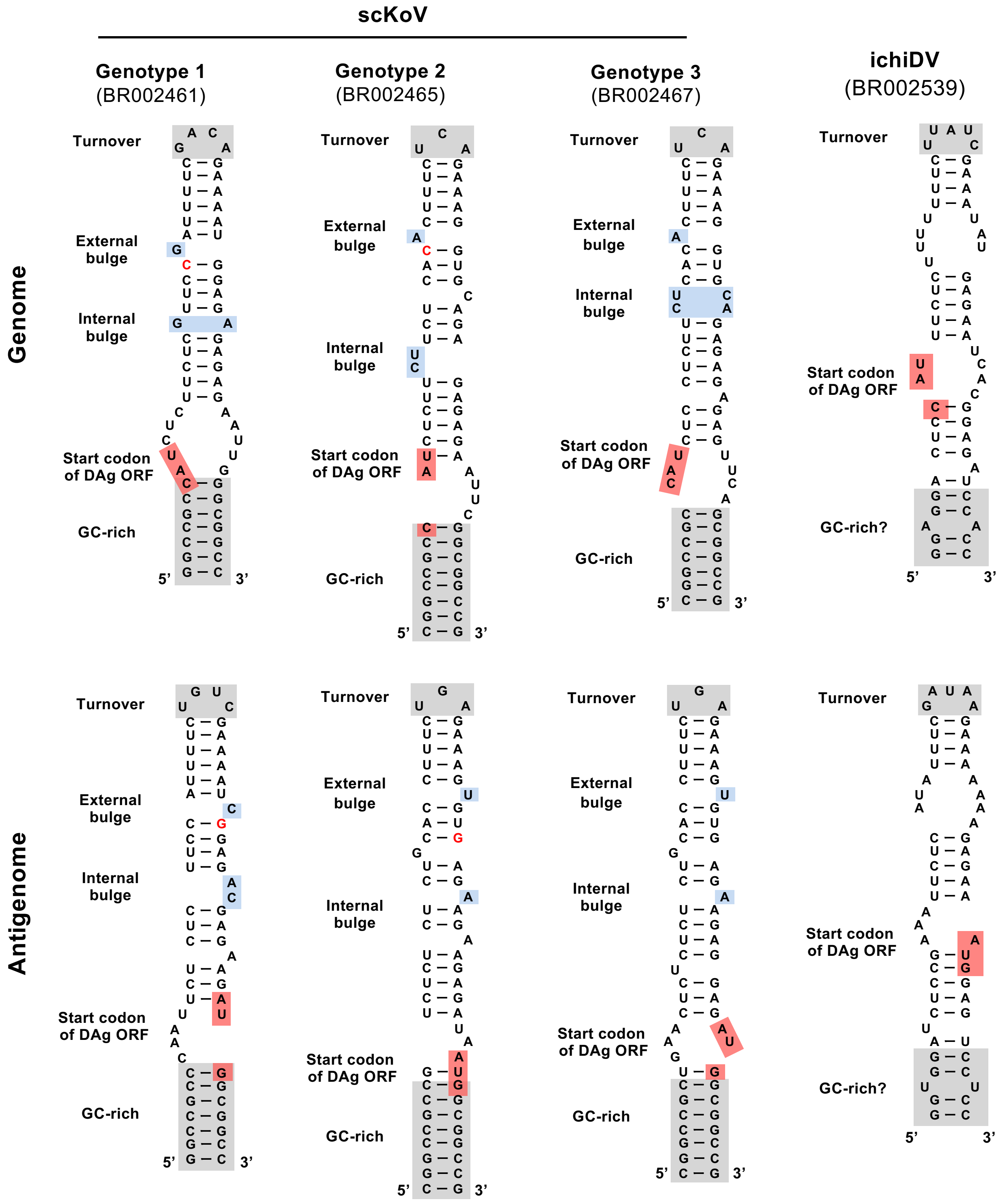


**Supplementary Figure 6. Secondary structures of putative promoter regions of scKoV and ichiDV.**

Secondary structures at the edge of the rod-like genomic and antigenomic RNAs upstream of the DAg ORF were predicted using the Mfold web server (Zuker et al., 2003). Conserved structural elements were annotated based on previous reports (Beard et al., 1996).

### Fig. S7

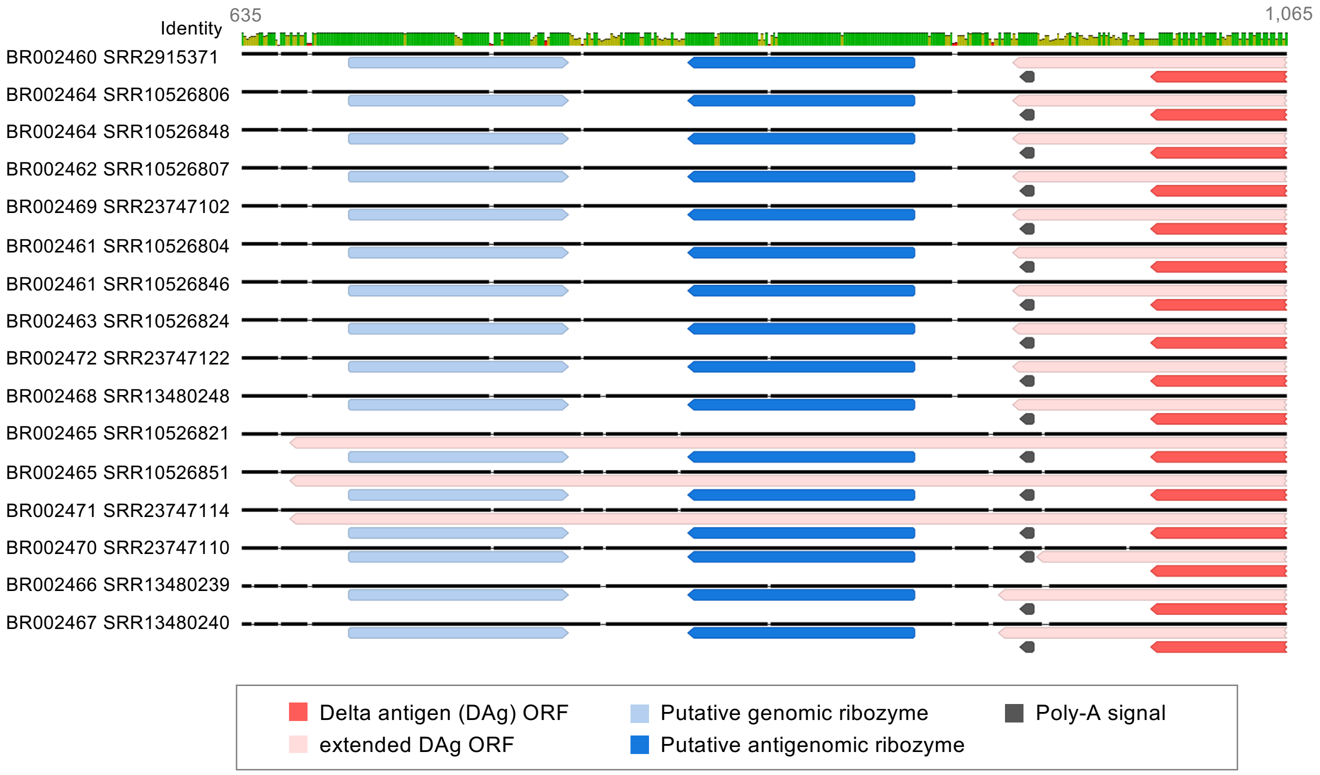


**Supplementary Figure 7. Comparison of extended DAg ORF of scKoVs.**
The complete nucleotide sequences of scKoVs were aligned using MAFFT v7.490 with E-INS-i algorithms (Katoh et al. 2013). An extended DAg ORF was predicted by replacing the original DAg stop codon with a sense codon. Colored arrows indicate annotations (ORFs, putative ribozymes, and poly-A signals). The numbers indicate nucleotide positions of the alignment.

Fig. S8

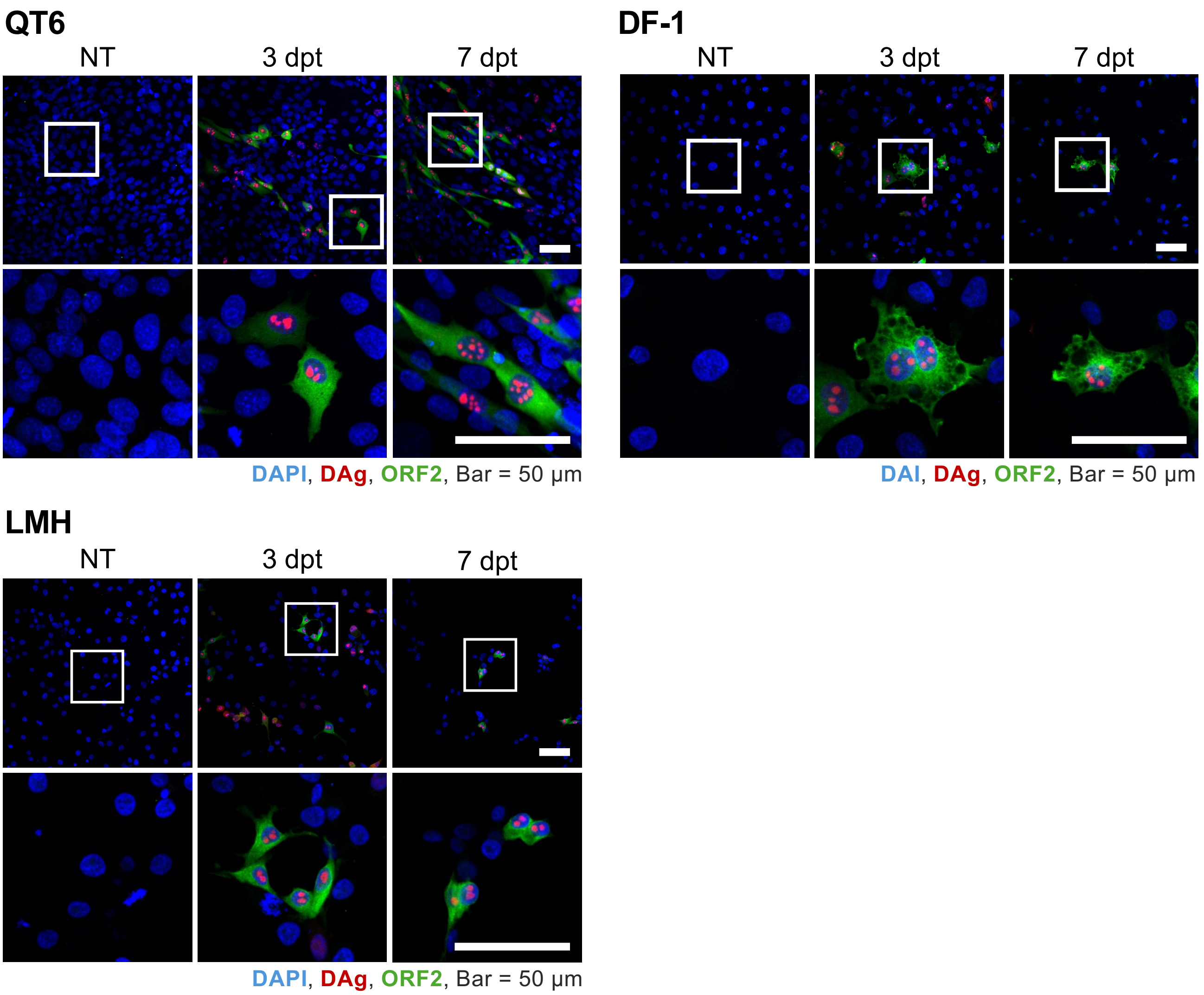


**Supplementary Figure 8. scKoV replication in avian cells.**
Indirect immunofluorescence analysis (IFA) of avian cells transfected with plasmids encoding the scKoV genome. Cells were visualized by fluorescence microscopy. QT6; a quail fibroblast cell line isolated from fibrosarcoma, DF-1; a chicken fibroblast cell line isolated from embryo; LMH; a chicken epithelial cell line isolated from hepatocellular carcinoma. Blue; DAPI, Red; DAg, Green; ORF2. Scale bars, 50 μm.

### Fig. S9

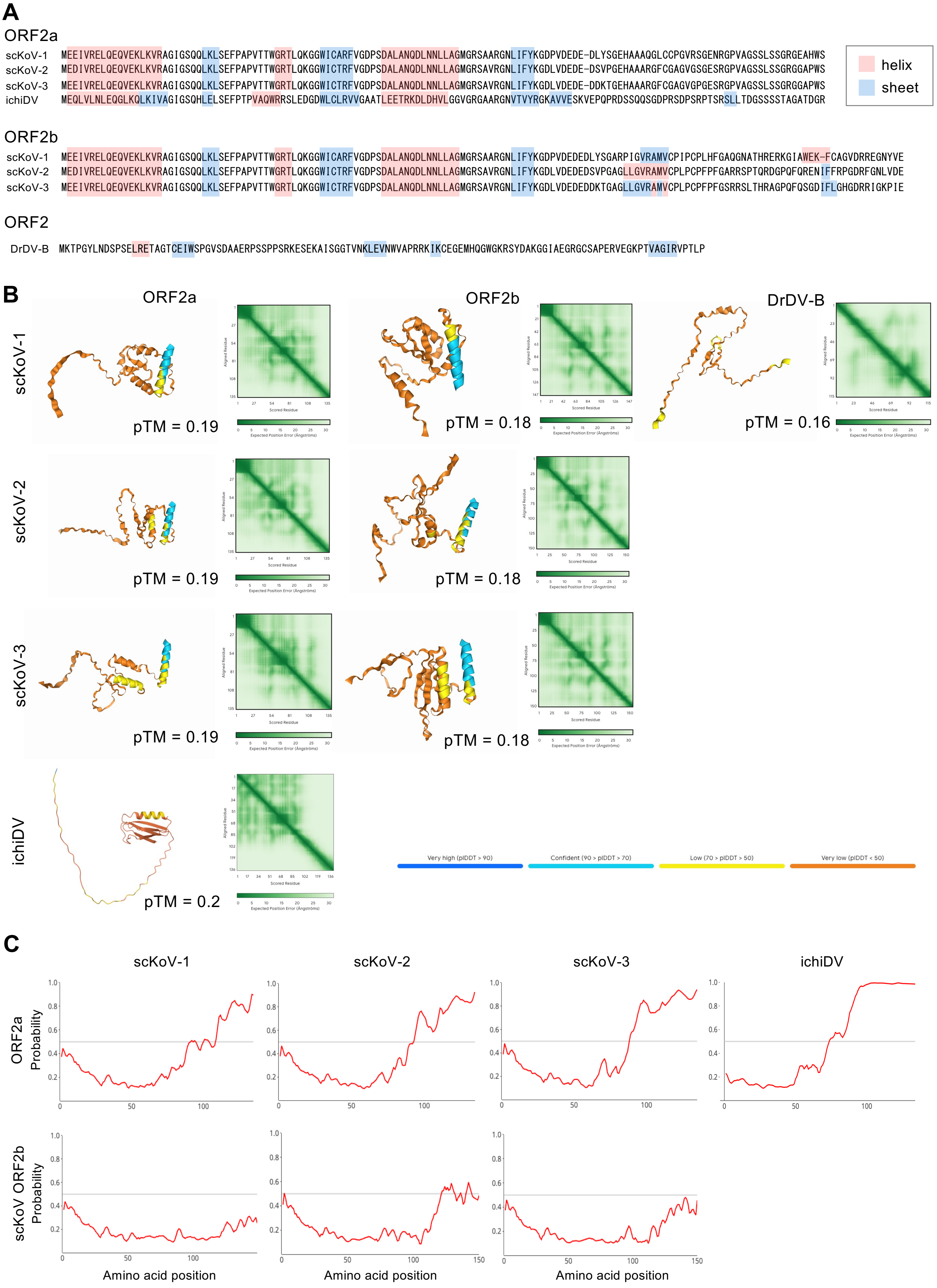
A
ORF2a
scKoV-1 MEEIVRELQEQVEKLKVRAGIGSQQLKLSEFPAPVTTWGRTLQKGGWICARFVGDPSDALANQDLNNLLAGMGRSAARGNLIFYKGDPVDEDE-DLYSGEHAAAQGLCCPGVRSGENRGPVAGSSLSSGRGEAHWS
scKoV-2 MEDIVRELQEQVEKLKVRAGIGSQQLKLSEFPAPVTTWGRTLQKGGWICTRFVGDPSDALANQDLNNLLAGMGRSAVRGNLIFYKGDLVDEDE-DSVPGEHAAARGFCGAGVGSGESRGPVAGSSLSSGRGGAPWS
scKoV-3 MEEIVRELQEQVEKLKVRAGIGSQQLKLSEFPAPVTTWGRTLQKGGWICTRFVGDPSDALANQDLNNLLAGMGRSAVRGNLIFYKGDLVDEDE-DDKTGEHAAARGFCGAGVGPGESRGPVAGSSLSSGRGGAPWS
ichiDV MEQLVLNLEQGLKQLKIVAGIGSQHLELSEFPTPVAQWRRSLEDGDWLCLRVVGAATLEETRKDLDHVLGGVGRGAARGNVTVYRGKAVVESKVEPQPRDSSQQSGDPRSDPSRPTSRSLLTDGSSSSTAGATDGR
helix
sheet
ORF2b
scKoV-1 MEEIVRELQEQVEKLKVRAGIGSQQLKLSEFPAPVTTWGRTLQKGGWICARFVGDPSDALANQDLNNLLAGMGRSAARGNLIFYKGDPVDEDEDLYSGARPIGVRAMVCPIPCPLHFGAQGNATHRERKGIAWEK-FCAGVDRREGNYVE
scKoV-2 MEDIVRELQEQVEKLKVRAGIGSQQLKLSEFPAPVTTWGRTLQKGGWICTRFVGDPSDALANQDLNNLLAGMGRSAVRGNLIFYKGDLVDEDEDSVPGAGLLGVRAMVCPLPCPFPFGARRSPTQRDGPQFQRENIFFRPGDRFGNLVDE
scKoV-3 MEEIVRELQEQVEKLKVRAGIGSQQLKLSEFPAPVTTWGRTLQKGGWICTRFVGDPSDALANQDLNNLLAGMGRSAVRGNLIFYKGDLVDEDEDDKTGAGLLGVRAMVCPLPCPFPFGSRRSLTHRAGPQFQSGDIFLGHGDRRIGKPIE
ORF2
DrDV-B MKTPGYLNDSPSELRETAGTCEIWSPGVSDAAERPSSPPSRKESEKAISGGTVNKLEVNWVAPRRKIKCEGEMHQGWGKRSYDAKGGIAEGRGCSAPERVEGKPTVAGIRVPTLP
B
ORF2a
ORF2b
DrDV-B
scKoV-1
pTM = 0.19
pTM = 0.18
pTM = 0.16
scKoV-2
pTM = 0.19
pTM = 0.18
scKoV-3
pTM = 0.19
pTM = 0.18
ichiDV
pTM = 0.2
Very high (pIDDT > 90)
Confident (90 > pIDDT > 70)
Low (70 > pIDDT > 50)
Very low (pIDDT < 50)
C
scKoV-1
scKoV-2
scKoV-3
ichiDV
ORF2a Probability
scKoV ORF2b Probability
Amino acid position
Amino acid position
Amino acid position

### Fig. S9

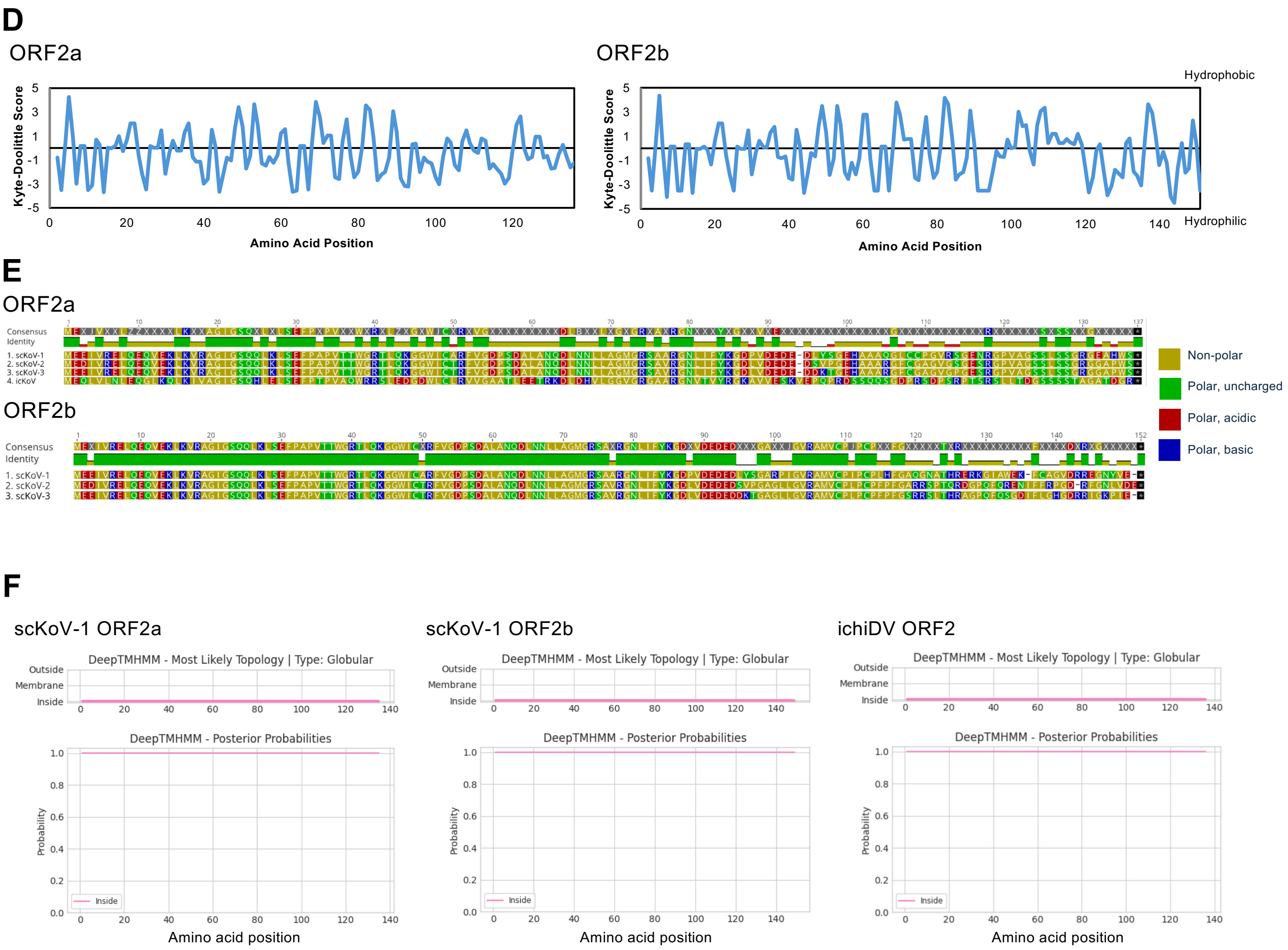


**Supplementary Figure 9. Characterization of ORF2 proteins of scKoV, ichiDV, and DrDV-B.** (**A**) The secondary structures of ORF2 proteins (ORF2a, unspliced isoform; ORF2b, spliced isoform) were predicted using JPred v4 (Drozdetskiy et al. 2015). Results of Jnet prediction were visualized in the alignments. (**B**) The 3D structures of ORF2 proteins were predicted using AlphaFold 3 (Abramson et al. 2024). (**C**) Predicted intrinsically disordered regions using AIUPred web server (Erdos and Dosztanyi et al. 2024). (**D**) The hydrophobicity value of each position of the amino acid alignments. (**E**) The amino acid sequence alignment of the ORF2 proteins was colored according to the polarity of each amino acid residue. (**F**) Predicted topology of the ORF2 proteins using DeepTMHMM v1.0.13 (Hallgren et al. 2022). The following sequences were used as representatives: BR002461 for scKoV genotype 1, BR002465 for genotype 2, BR002467 for genotype 3, BR002539 for ichiDV, and MT649206 for DrDV-B.

Fig. S10

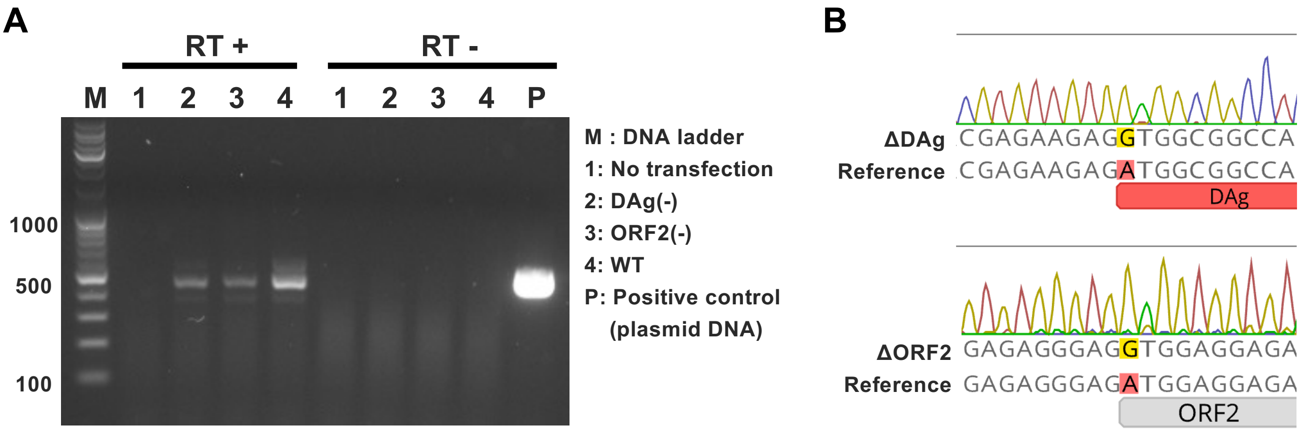


**Supplementary Figure 10. Confirmation of introduced mutations in translation-deficient mutants (scKoV-ΔDAg and scKoV-ΔORF2).**
To verify the persistence of the introduced mutations in scKoV-ΔDAg and scKoV-ΔORF2, RNA was extracted from QT6 cells 7 days post-transfection and used for sequencing of the target regions. (**A**) Endpoint RT-PCR of the target region. To exclude plasmid DNA contamination, RT-PCR and control PCR (without reverse transcription) were performed. Amplification products of the expected size were observed only in the RT-PCR condition. (**B**) Sanger sequencing of the RT-PCR products confirmed the presence of the introduced mutations at 7 days post-transfection.

Fig. S11

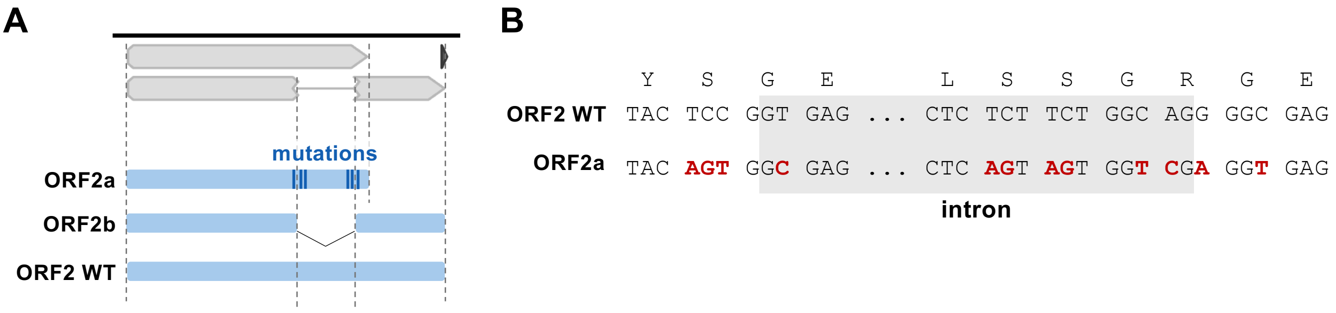


**Supplementary Figure 11. Plasmid constructs used for expression of scKoV ORF2 isoforms.** (**A**) Schematic diagram of plasmid inserts expressing ORF2a, ORF2b, and ORF2 wild type (WT). (**B**) Mutations introduced into the plasmid expressing ORF2a. Introduced mutations are highlighted in red.

### Fig. S12

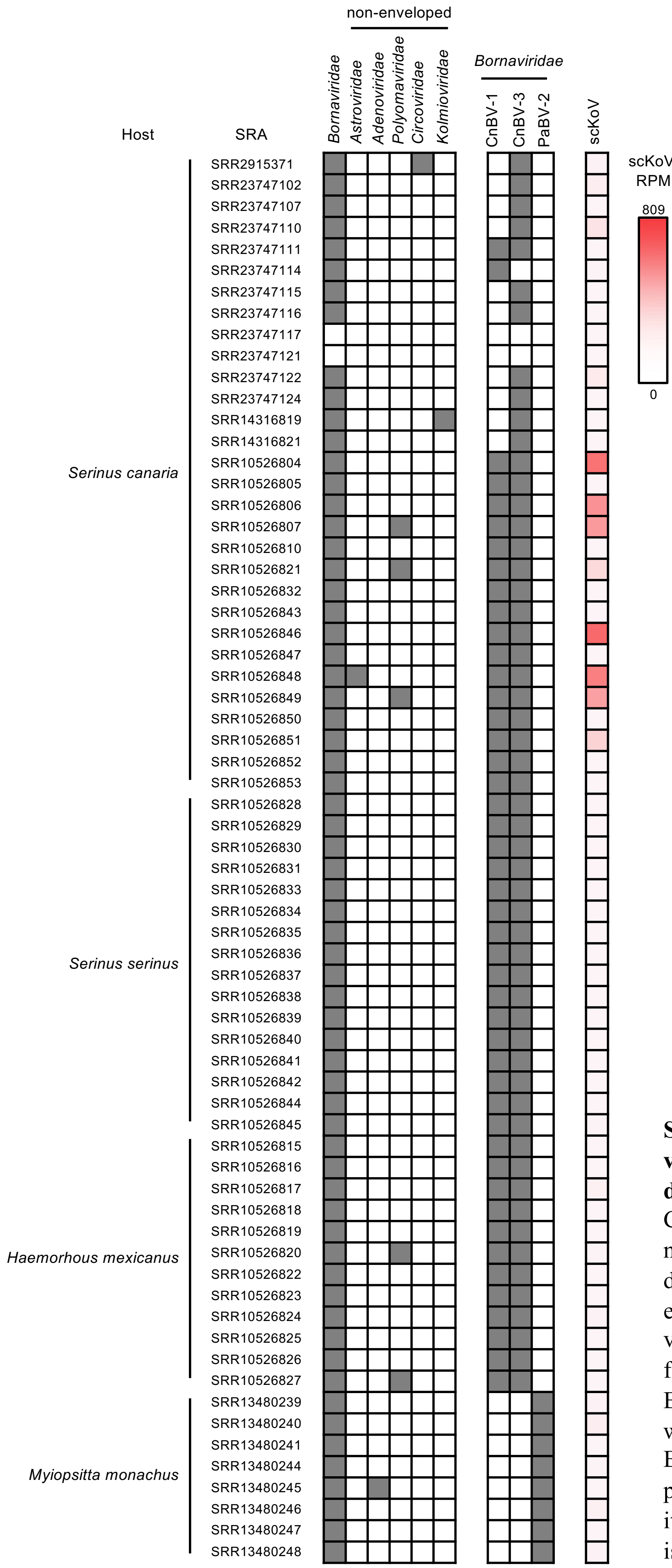


**Supplementary Figure 12. Co-existing viruses in scKoV-positive public RNA-seq datasets (full version).**
Co-existing viruses and scKoV-derived read number detected in each public RNA-seq dataset are shown. All co-existing viruses, excluding retroviruses and environmental viruses, were identified in contigs assembled from scKoV-positive RNA-seq datasets using BLASTx (left panel). Contigs of *Bornaviridae* were further classified into each virus using BLASTn (middle panel). Black indicates the presence of a viral contig, while white indicates its absence. Read per million (RPM) of scKoV is shown as a heatmap (right panel).

### Fig. S13

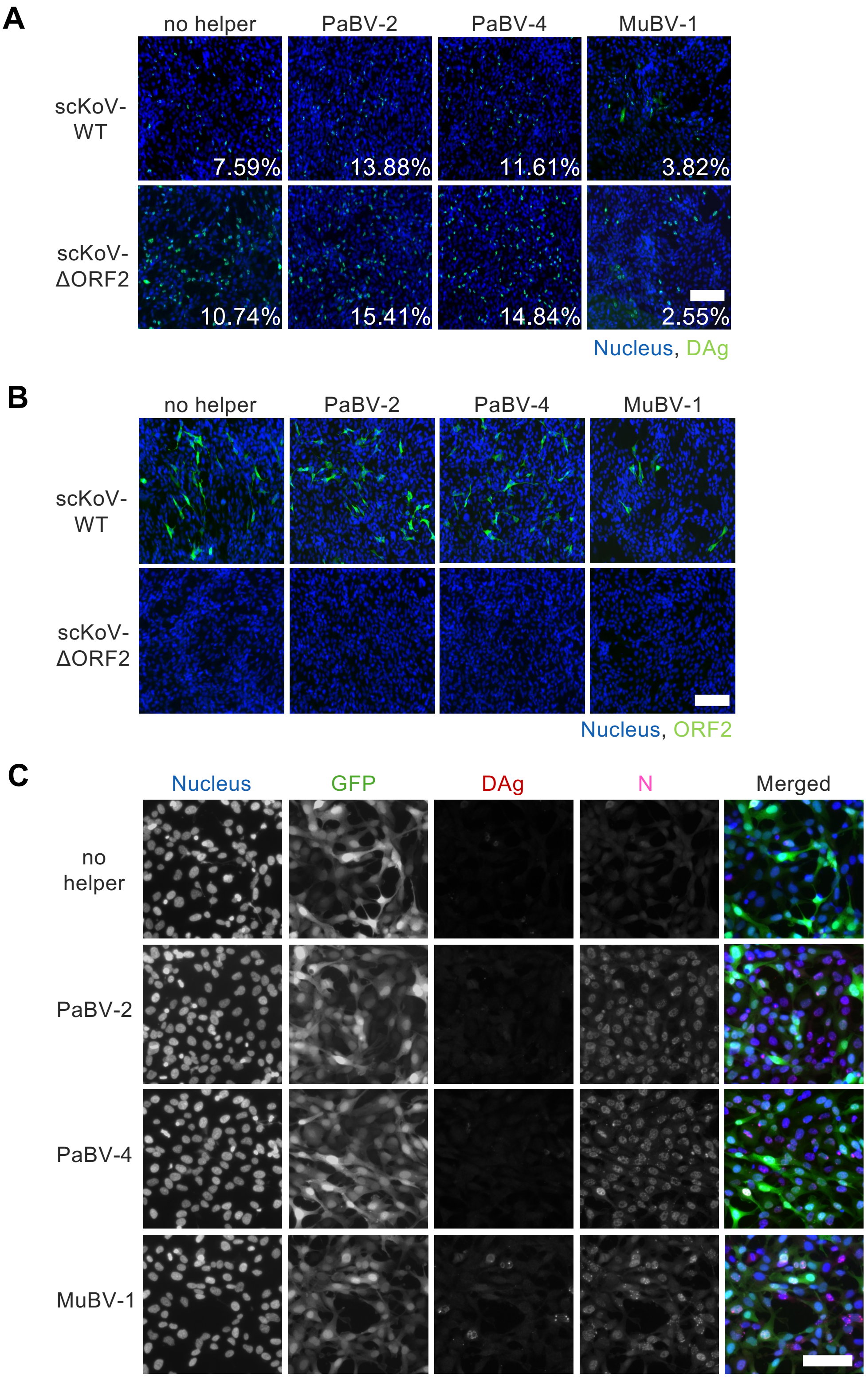


**Supplementary Figure 13. scKoV transmission in the presence of avian bornaviruses.**
(**A and B**) Immunofluorescence images of producer QT6 cells infected with scKoV in the presence or absence of avian bornaviruses including parrot bornavirus 2 (PaBV-2), parrot bornavirus 4 (PaBV-4), and munia bornavirus 1 (MuBV-1). Numbers in the lower right corners indicate the percentage of scKoV-positive cells. Blue, DAPI; green, DAg (A) or ORF2 (B). Scale bars, 100 µm. (**C**) Representative immunofluorescence images of recipient cells after transmission of scKoV-ΔORF2, acquired by fluorescence microscopy. Blue, DAPI; green, GFP; red, scKoV DAg; magenta, bornavirus N protein. Scale bar, 50 µm.

### Fig. S14

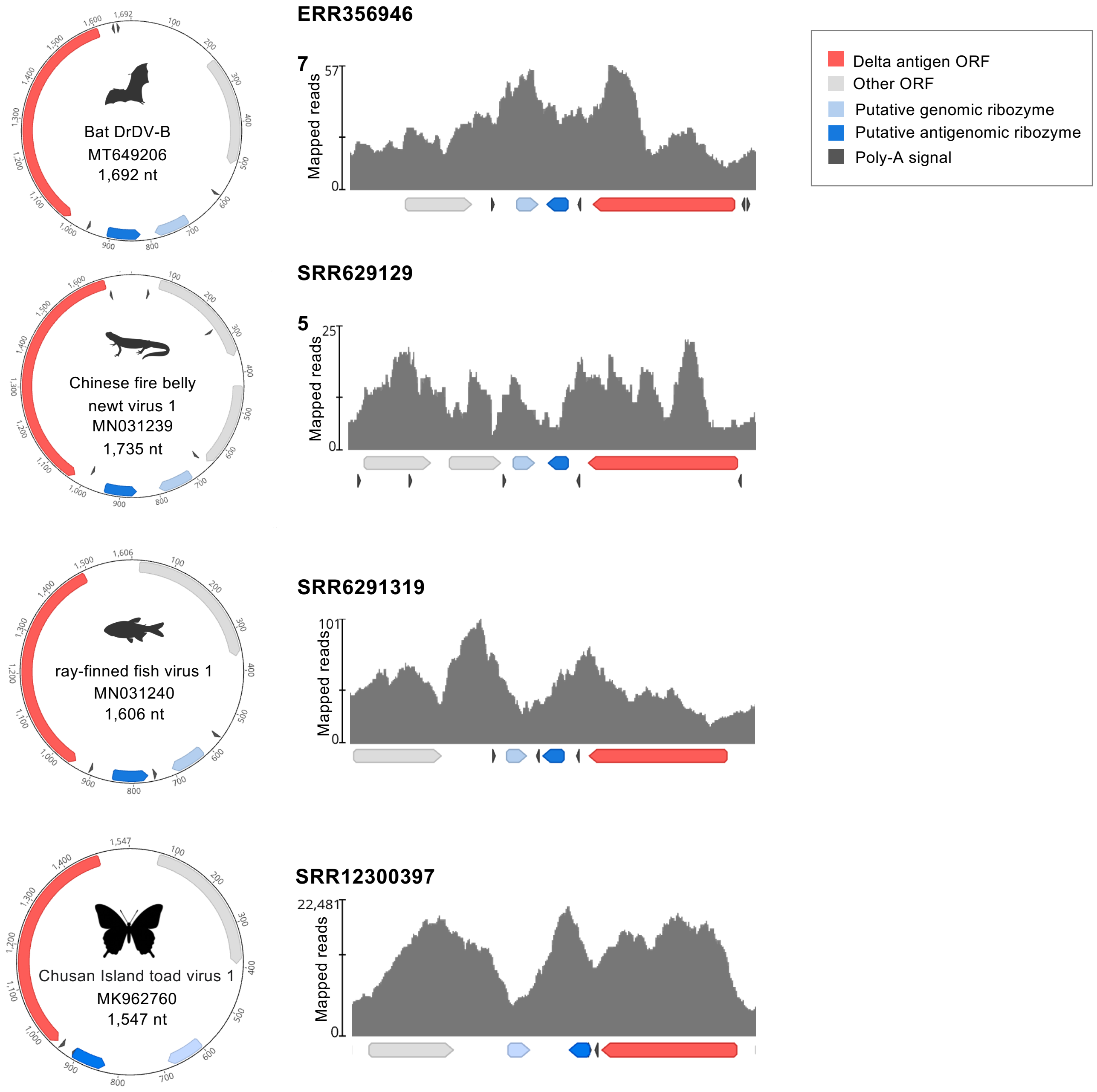


**Supplementary Figure 14. Genome structure (left panel) and transcriptional pattern (right panel) of KoVs with genomic poly-A signals.**

Colored arrows indicate annotations (ORFs, putative ribozymes and poly-A signals). The numbers indicate nucleotide positions. For Babylonia areolata KoV, no obvious HDV-like or hammerhead ribozymes were predicted. Illustrations were created with BioRender.com or downloaded from PhyloPic (https://www.phylopic.org).

Fig. S15

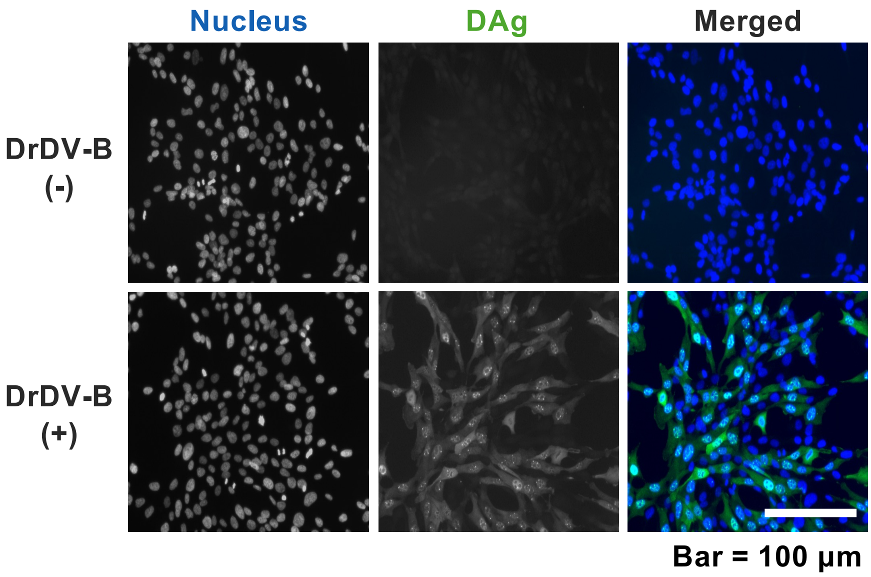


**Supplementary Figure 15. Rescue of DrDV-B.**
A plasmid encoding a 1.2× length DrDV-B genome was constructed and transfected into QT6 cells. At 7 days post-transfection (dpt), cells were examined by indirect immunofluorescence assay (IFA) using fluorescence microscopy. DrDV-B DAg was detected using rabbit antiserum against DrDV-B DAg. Blue, DAPI; green, DAg. Scale bar, 100 μm.
